# Interferon response and epigenetic modulation by *SMARCA4* mutations drive ovarian tumor immunogenicity

**DOI:** 10.1101/2023.08.08.552544

**Authors:** Melica Nourmoussavi Brodeur, Higinio Dopeso, Yingjie Zhu, Ana Leda F. Longhini, Andrea Gazzo, Siyu Sun, Richard Koche, Rui Qu, Pierre-Jacques Hamard, Yonina Bykov, Hunter Green, Katherine B. Chiappinelli, Melih Arda Ozsoy, Thais Basili, Rui Gardner, Sven Walderich, Elisa DeStanchina, Benjamin Greenbaum, Mithat Gönen, Britta Weigelt, Dmitriy Zamarin

**Author notes:** Corresponding author: Dmitriy Zamarin, MD PhD, Gynecologic Medical Oncology, Department of Medicine, Memorial Sloan Kettering Cancer Center, 1275 York Avenue, New York, NY 10065, USA.

## Abstract

Cell-intrinsic mechanisms of immunogenicity in ovarian cancer (OC) are not well understood. The presence of damaging mutations in the SWI/SNF chromatin remodeling complex, such as the *SMARCA4* (BRG1) catalytic subunit, has been associated with improved response to ICB, however the mechanism by which this occurs is unclear. The aim of this current study was to examine the alterations in tumor cell-intrinsic and extrinsic immune signaling caused by *SMARCA4* loss. Using OC models with loss-of-function mutations in *SMARCA4*, we found that *SMARCA4* loss resulted in increased cancer cell-intrinsic immunogenicity, characterized by upregulation of long-terminal RNA repeats such as endogenous retroviruses, increased expression of interferon-stimulated genes, and upregulation of antigen presentation machinery. Notably, this response was dependent on IRF3 signaling, but was independent of the type I interferon receptor. Mice inoculated with cancer cells bearing *SMARCA4* loss demonstrated increased activation of cytotoxic T cells and NK cells in the tumor microenvironment as well as increased infiltration with activated dendritic cells. These results were recapitulated when animals bearing *SMARCA4-*proficient tumors were treated with a BRG1 inhibitor, suggesting that modulation of chromatin remodeling through targeting *SMARCA4* may serve as a strategy to reverse immune evasion in OC.

## Introduction

Immune targeted therapies, such as immune checkpoint blockade (ICB), have shown dramatic success in multiple cancer types, including some gynecologic cancers like endometrial^1,2^ and cervical cancers^3^. To date, ICB trials in patients with ovarian cancer (OC) have yielded disappointing response rates with no clear biomarker for response^4^. Nonetheless, several studies have demonstrated that a subset of OCs is characterized by an immunogenic tumor microenvironment (TME) that could make them sensitive to ICB^4,5^.

As OC is the most lethal gynecologic cancer^6^, a better understanding of the tumor cell-intrinsic mechanisms mediating immune response and resistance to immune cell killing is needed. In several cancer types, epigenetic modifications have been linked to changes in the immune TME and increased clinical benefit of ICB^7–10^. Many of these epigenetic changes are directly or indirectly linked to damaging mutations in genes encoding subunits of the SWItch/Sucrose Non-Fermentable (SWI/SNF) chromatin remodelling complex^11–13^. This complex normally enables selective gene expression and DNA repair through modulation of DNA compaction and accessibility^14^. With over 20% of human cancers harboring mutations in SWI/SNF genes, its implication in cancer development through dysregulation of oncogenes and tumor suppressor genes has started to be uncovered^14^. For example, *SMARCB1*-mutated rhabdoid tumors have been shown to exhibit increased immune cell infiltration and immune cell killing function^11^. Furthermore, a clinical study evaluating ICB (anti-PDL1) response in lung cancers showed increased response in *SMARCA4*-mutated compared to *SMARCA4*-wildtype tumors^13^. We and others have demonstrated similar immune changes in the TME of SWI/SNF-deficient OCs^5,15^, however the mechanism by which this immune response occurs are still not well understood. In particular, a subset of aggressive rhabdoid-like OCs, called small cell carcinoma of hypercalcemic type (SCCOHT), harbor inactivating mutations in the catalytic subunit of the SWI/SNF complex called *SMARCA4* (protein product known as BRG1)^5^. Despite a low tumor mutational burden (TMB), one of the known predictive biomarkers of ICB response^16^, these *SMARCA4*-mutated tumors have prominent T cell infiltration and PDL1-expressing tumor-associated macrophage,s and case reports show clinical responses to ICB^5^. The mechanisms underlying this immunogenicity remain unclear.

Here, we sought to examine the alterations in tumor cell-intrinsic and extrinsic immune signaling caused by *SMARCA4* loss using syngeneic OC murine models. We demonstrate that the underlying immunogenicity in *SMARCA4*-mutated OCs is driven by upregulation of downstream type I interferon (IFN)-stimulated genes (ISGs) and MHC class I expression. In animal models, this led to delayed tumor growth and was associated with increased tumor infiltration of dendritic cells and activated T cells. Treatment of *SMARCA4* wild-type cells/tumors with BRG1-targeted inhibitors and degraders resulted in activation of type I IFN signaling in cancer cells, decreased tumor burden, and enhanced tumor immune infiltration. Our findings generate rationale for exploration of alterations in SWI/SNF-mediated signaling as biomarkers of response to immunotherapy and for targeting of BRG1 as a potential strategy to enhance tumor cell immunogenicity as a therapeutic strategy.

## Materials and Methods

### Cell lines

The ID8-Luciferase (ID8) cell line was kindly provided by Dr. Renier Brentjens’ lab. The UPK10- GFP (UPK10) cell line was a kind gift from Dr. Jose Conejo-Garcia’s lab. OAW28 cells were purchased commercially from the European Collection of Authenticated Cell Cultures (85101601-1VL, Sigma). B16-F10 cells were originally obtained from the American Type Culture Collection (ATCC, #CRL-6475). All cell lines (**Table S1**) underwent routine authentication using short tandem repeat (STR) profiling and were tested for mycoplasma using the PCR-based Universal Mycoplasma Detection kit (#30-1012K, ATCC). ID8 and UPK10 were cultured in RPMI medium supplemented with 10% fetal bovine serum (FBS), 1% penicillin/streptomycin. B16-F10 cells were maintained in RPMI medium supplemented with 7.5% FBS, 1% penicillin/streptomycin. OAW28 cell line was maintained in DME, 2 mM glutamine, 1mM sodium pyruvate, 20 IU/L bovine insulin with 10% FBS, 1% penicillin/streptomycin. Cells were maintained in a 5% CO_2_ atmosphere at 37°C. Experiments were carried out with cells between passage 2 to 10 (at 80% confluence).

Isogenic *SMARCA4* knockout (KO) murine (ID8, UPK10) OC cell lines were generated using CRISPR-Cas9 lentiviral transduction and puromycin selection (**Fig S1A**). Isogenic murine cell lines further underwent clonal selection using single-cell sorting (**Fig S1B**). A panel of 5 human OC cell lines were tested for *SMARCA4* baseline expression to identify the cell line with highest expression (**Fig S1C**). Isogenic single guide *SMARCA4* (sg*SMARCA4*) human OAW28 cell lines were generated using CRISPR-Cas9 lentiviral transduction and puromycin selection (**Fig S1D**).

### SMARCA4-knockout cell line establishment

Stable sg*SMARCA4* cell lines were created with ID8, UPK10 and OAW28 models. Individual gRNAs (**Table S2**) against murine and human *SMARCA4* were cloned into pLentivectorCRISPR v2 (52961, Addgene). Non-target control (NTC) was used as control. Lentiviral particles were produced at Memorial Sloan Kettering Cancer Center’s (MSK) Gene Editing and Screening Core Facility. Lentiviral titers (TU/mL) were first determined (see section *Viral Titration*). Each cell line was infected for 24h in the presence of 10ug/mL polybrene (TR-1003-G, Millipore) and then selected for 14 days in puromycin (2 µg/mL for ID8 and 8ug/mL for UPK10; ThermoFisher Scientific). Transduction efficiency was confirmed by quantitative reverse transcription polymerase chain reaction (qRT-PCR) and western blot. To generate single cell clones for ID8 and UPK10 lines, single-cell suspensions stained with DAPI were sorted by Aria cell sorter to distribute cells into the wells of a 96-well plate with 100uL of culture medium. Each single-cell clone was expanded, then tested for *SMARCA4*-mutation using Sanger sequencing and validated using immunofluorescence.

For generation of stable sg*SMARCA4* B16-F10 cell line, individual gRNAs (**Table S2**) against murine *SMARCA4* encoded in the px459 vector were generated by the MSK Gene Editing and Screening Core Facility. Cas9-expression empty vector (EV) was used as a control. Cells were transfected with all three gRNAs by electroporation using the Eppendorf Multiporator. Cells were allowed to recover for 48 hours, and then selected using 3 µg/mL puromycin for 14 days. Single cell clones were generated by seeding a 96-well plate with a limiting dilution of the single cell suspension. Each single-cell clone was expanded, then tested for *SMARCA4* mutation using Sanger sequencing and validated using western blot.

### Sanger sequencing

Total DNA was extracted from isogenic cell lines using the DNeasy Blood & Tissue Kit (69504, Qiagen), according to manufacturer’s instructions. DNA was amplified using AmpliTaq Gold 360 Master Mix (4398886, Thermofisher Scientific) and primers (**Table S3**). PCR products were purified using ExoSAP-IT PCR product Cleanup (78200, Thermofisher Scientific) and subjected to Sanger sequencing (Genewiz) using the same primers. Sequencing traces were reviewed using SnapGene (http://www.snapgene.com) to assess the sgRNA editing.

### Lentiviral transduction for doxycycline-inducible knockdowns

MAVS and IRF3 knockdown cell lines were created in the murine model ID8. Short hairpin RNAs (shRNAs) were designed using the SplashRNA algorithm^17^. A previously described optimized lentiviral miR-E expression backbone system was used for constitutive SREP (red, puromycin) and inducible LT3RENIR (red, Neomycin) expression^18^. Individual LT3RENIR lentiviral miR-E-based expression vectors containing shRNA against MAVS, IRF3 and NTC (**Table S4**) were cloned, and lentiviral particles were produced in MSK’s Gene Editing and Screening Core Facility. Lentiviral titers (TU/mL) were first determined (see section *Viral Titration*). ID8 cells were infected for 24h and then selected for 14 days in geneticin (50mg/mL; 10131035. ThermoFisher Scientific). shRNAs were induced with 6 days of doxycycline treatment prior to experiments. Knockdown efficiency was confirmed by qRT-PCR and western blot. The short hairpin sequences used are described in **Table S4**.

### Viral titration

Cells were seeded in 24-well plates (25,000-50,000 cells/well). On the following day, the cells were transduced for 6h with 5-fold serial dilutions (protocol 25–78125, Horizon Discovery) of lentivirus stock. Cells were incubated for 48h. Transduction efficiency was confirmed by the number of puromycin-or geneticin-selected colonies.

### RNA-sequencing analyses

Total RNA extracted from ID8 and OAW28 cells with or without *SMARCA4*-deficiency using the RNeasy Mini Kit (74104, Qiagen) was subjected to ribodeplete RNA-sequencing at MSK’s Integrated Genomics Operation (IGO). For coding gene mapping and analysis, RNA-sequencing reads were aligned to the reference mouse (version mm10) and human (version hg38) genomes using STAR aligner^19^, and the number of reads counted were mapped to each gene using GenomicAlignments package in Bioconductor^20,21^. Expression fold changes between sg*SMARCA4* and sgNTC cells were calculated from DESeq2^22^, which was performed using raw read counts. P-values obtained from DESeq2 were corrected for multiple testing using the Benjamini-Hochberg method^23^. ClusterProfiler^24^ was employed for gene set enrichment analysis (GSEA) according to fold-change and differentially expressed genes. Read counts were normalized by DESeq2 for visualization.

For non-coding gene mapping, counts filtering and normalization and differential expression analysis, RNA-sequencing reads were 3’ trimmed for base quality less than 20 using Skewer^25^. Reads shorter than 40 after trimming were removed. The quality score of the remaining bases were sorted, and the quality at the 20th percentile was computed. Reads with quality at the 20th percentile less than 15 were discarded. Only paired reads which both passed the quality check were mapped to the reference genome (mm10 or hg38) using STAR v2.7 with default parameters^19^. Gene counts were assigned based on Gencode annotation using featureCounts (subread-1.4) with the external Ensembl annotation. Repeats counts per element family were primarily quantified against RepeatMasker using featureCounts (subread-1.4) and then adding the counts of the unassigned reads that mapped to Repbase consensus sequence. Repeat counts of a given family is the sum of mapped reads to RepeatMasker and unmapped reads against Repbase.

Expression of repeats and coding genes was normalized across samples using trimmed-mean of M-values (TMM) in edgeR. The size factor for each sample was calculated using calcNormFactors based on coding genes alone as described in a previous study^26^. Low-count genes/repeats in each sample with counts per million (CPM) smaller than 2 were removed before calculating the size factor. Log2 transformed CPM were used for downstream visualization.

Differential expressions of coding genes and repetitive elements were analyzed separately using DESeq2 (v1.33.4)^22^.

### Assay for transposase-accessible chromatin (ATAC) sequencing and epigenome analyses

Freshly harvested ID8 sg*SMARCA4* and sgNTC cells were directly sent to MSK’s Epigenetics Research Innovation Lab. ATAC was performed as previously described^27^ using the Tagment DNA TDE1 Enzyme (Illumina, 20034198). Sequencing libraries were prepared using the ThruPLEX DNA-Seq Kit (Takarabio, R400676) and sent to the MSK IGO core facility for sequencing on a NovaSeq 6000. Raw sequencing reads were trimmed and filtered for quality (Q>15) and adapter content using version 0.4.5 of TrimGalore (https://www.bioinformatics.babraham.ac.uk/projects/trim_galore) and running version 1.15 of cutadapt and version 0.11.5 of FastQC. Version 2.3.4.1 of bowtie2 (http://bowtie-bio.sourceforge.net/bowtie2/index.shtml) was employed to align reads to mouse assembly mm10 and alignments were deduplicated using MarkDuplicates in Picard Tools v2.16.0. Enriched regions were discovered using MACS2 (https://github.com/taoliu/MACS) with a p-value setting of 0.001, filtered for blacklisted regions (http://mitra.stanford.edu/kundaje/akundaje/release/blacklists/mm10-mouse/mm10.blacklist.bed.gz), and a peak atlas was created using +/- 250 base pairs around peak summits. The BEDTools suite (http://bedtools.readthedocs.io) was used to create normalized bigwig files. Version 1.6.1 of featureCounts (http://subread.sourceforge.net) was used to build a raw counts matrix and DESeq2 was employed to calculate differential enrichment for all pairwise contrasts. Peak-gene associations were created by assigning all intragenic peaks to that gene, while intergenic peaks were assigned using linear genomic distance to transcription start sites (TSS). Pathway enrichment was calculated by assigning each gene a unique score based on the associated peak with the greatest magnitude change and running GSEA in pre-ranked mode. Motif signatures were obtained using Homer v4.5 (http://homer.ucsd.edu) on differentially enriched peak regions. IRF3-enriched regions were downloaded from the Peak Browser of the mm10 version of the ChIP-Atlas database (http://chip-atlas.org) and displayed using Integrative Genomics Viewer (IGV)^28^.

### Transcript abundance (qRT-PCR)

Total RNA (500-1000ng) extracted from isogenic cell lines was used to generate cDNA with the Superscript VILO Master Mix (11755050, Thermofisher Scientific) according to manufacturer’s instructions. In murine and human cells, qRT-PCR (**Table S5**) was performed using TaqMan Universal Master Mix II, with UNG (4440044) for IFN genes and PowerUp SYBR Green (A25776, Thermofisher Scientific) for endogenous retroviral genes using StepOnePlus Real-Time PCR System (Applied Biosystems, ThermoFisher). β-actin was used as the reference gene. The ΔΔCT method was used to calculate relative expression levels, as previously described^29^. All qRT-PCR assays were carried out in triplicate and then repeated with new cDNA synthesis. Reverse transcription-negative cDNA synthesis reactions were performed for each gene tested.

### Treatments

For BRG1/BRM-targeted treatment, ID8 and UPK10 parental cells were seeded in a 6-well plate at 50,000 cells/well. After 24h, cells were treated for 48h with 1 µM of the dual BRG1/BRM inhibitor (#HY-119374, MedChemExpress) or dual BRG1/BRM proteolysis-targeting chimera (PRTD-0043, Prelude Therapeutics) or vehicle (DMSO). Cells were then collected after 48h for RNA extraction and qRT-PCR of ISGs.

To validate the efficiency of IFNAR blockade, ID8 and OAW28 NTC cells were seeded in a 6-well plate at 50,000-200,000 cells/well. After 24h, cells were treated for 24h with or without recombinant murine IFNα (12100-1, PBL Assay Science) or human IFNα (11100-1, PBL Assay Science) at 2000U/mL. After 24h, cells were treated for 48h with mock (media) or anti-human IFNAR2 (21385-1, PBL Assay Science) or anti-mouse IFNAR-1 (BE0241, BioXCell) at 8-10 ug/mL. Cells were collected after 48h for RNA extraction and qRT-PCR of IFN genes.

### Protein analyses

Standard western blotting was conducted^30^. Primary antibodies against MAVS (BS-4053R, Bioss; 1:500), IRF3 (MA5-32348, Invitrogen; 1:500) and BRG1 (49360, Cell Signaling; 1:400) were used. Alpha-tubulin (2144S, Cell Signaling; 1:1000) and vinculin (sc-73614, Santa Cruz; 1:5000) were used as reference protein. Conjugated secondary anti-rabbit antibodies were used and detected using the iBright Systems (Invitrogen, FL1500). For detection of BRG1 and vinculin, HRP-linked secondary anti-rabbit (7074P2, Cell Signaling; 1:10000) and anti-mouse (7076P2, Cell Signaling; 1:10000) antibodies were used respectively. For all blots, quantification and analysis were performed using iBright Analysis Software (V5.1.0). Experiments were performed in triplicate.

For immunofluorescence analysis, cells were grown on glass coverslips and fixed with 4% formaldehyde (28908, ThermoFisher Scientific), blocked in 10% normal Goat Serum (Vector Laboratories, S-1000) and then incubated with primary antibody against BRG1 (3508, Cell Signaling; 1:200) or IRF3 (MA5-32348, Invitrogen; 1:100). The samples were incubated overnight at 4°C. Conjugated secondary antibody Alexa-488 goat anti-rabbit (A-11034, ThermoFisher Scientific; 1:500) was incubated for 2h. Slides were mounted using Gold Prolong antifade reagent containing DAPI (P36930, ThermoFisher Scientific). Slides were scanned at MSK’s Molecular Cytology Core using a Pannoramic Scanner (3DHistech, Budapest, Hungary) with a 40x/0.95NA air objective. No post-acquisition processing was performed besides minor adjustments of brightness and contrast, applied equally to all images, using the QuPath software^31^ (Version 0.3.2). Experiments for each condition were performed in triplicate.

### In vivo studies

Six to 10-week-old female C57BL/6J (RRID:IMSR_JAX:000664, The Jackson Laboratory) and B6(Cg)-*Tyr^c-2J^*/J (RRID:IMSR_JAX:000058, The Jackson Laboratory) mice were used for *in vivo* studies and were cared for in accordance with the guidelines approved by the MSK Institutional Animal Care and Use Committee (IACUC) and Research Animal Resource Center. ID8 sgNTC and ID8 sg*SMARCA4* tumors were generated by injecting 10 million cells with PBS intraperitoneally (IP). ID8 tumors were generated by injecting 2 million cells with PBS IP. Tumors were measured weekly using IVIS Spectrum *in vivo* imaging and tumor burden was calculated using the Live Imaging 4.0 software. Body weight and clinical signs were also assessed. At the end of the study, the ascites was collected for flow cytometry analysis.

B16-F10 sgEV and sg*SMARCA4* tumors were generated by injecting 150,000 cells in 100 µL PBS intradermally in the flank. Tumors were measured every 48h using a caliper and volume was calculated using the formula *volume = length * width * height * 0.5*. Animals were euthanized at the humane endpoint of 1000 mm^3^ tumor volume, weight loss of ≥20%, or acute physical distress in accordance with IACUC guidelines. At the end of the study, the tumors were collected for flow cytometry analysis.

### Flow cytometry

After tumor harvest, the tumors were chopped and digested using DNAse I (10104159001, Sigma) and Liberase (5401020001, Sigma) for 30 minutes at 37°C. Samples were strained through 70 µM filters and centrifuged. Percoll 40% was used to remove fatty cells in ID8 tumors. Viability stain (**Table S6**) was performed by incubating cells in PBS for 15 minutes. Cells were washed in PBS, incubated in Fc-blocking buffer 2.4G2 for 15 minutes and stained with the extracellular antibodies (**Table S6**) in Brilliant stain buffer (563794, BD biosciences) for ID8 tumors and FACS buffer for B16-F10 cells for 30 minutes on ice. Cells were fixed and washed using Transcription factor staining buffer set (00-5523-00, ThermoFisher Scientific) and permeabilization buffer according to manufacturer’s protocol, and then stained with the intracellular antibodies (**Table S6**) in permeabilization buffer for 30 minutes on ice. Cells were resuspended in 200uL of FACS buffer and acquired on the 5-laser Cytek Aurora spectral flow cytometer (Cytek Biosciences). Data was analyzed using FlowJo software (BD Biosciences).

To examine PD-L1 and MHC1 expression in ID8 and OAW28 sgNTC and sg*SMARCA4* cells, cultured isogenic cells were harvested with trypsin and quenched with media. Cell pellets were washed in PBS and collected by centrifugation. Viability stain (**Table S6**) was performed by incubating cells in PBS for 15 minutes. Cells were washed in PBS and incubated in Fc-blocking buffer 2.4G2 for 15 minutes. ID8 and OAW28 cells were stained with extracellular antibodies (**Table S6**) for 30 minutes in blocking buffer. Cells were resuspended in 200uL of FACS buffer and acquired on the 5-laser Cytek Aurora spectral flow cytometer (Cytek Biosciences). Data was analyzed using FlowJo software (BD Biosciences).

### Transcriptomic analysis of TCGA data

Normalized RNA-sequencing data from The Cancer Genome Atlas (TCGA) pan-cancer study^32^ were obtained (https://gdc.cancer.gov/about-data/publications/pancanatlas). Immune-related scores were obtained from Thorsson et al^33^. Tumor types with cases in both datasets with a tumor purity ≥20% were included to assess *SMARCA4* expression levels. To avoid confounding factors related to tissue type, *SMARCA4* groups in each cancer type were stratified: 1) 25% cases with lowest *SMARCA4* expression levels (*SMARCA4*-low); 2) 25% cases with highest *SMARCA4* expression levels (*SMARCA4*-high). Tumor type with <3 cases in either group were excluded from our analyses. Then, *SMARCA4*-low expressing cases were compared with *SMARCA4*-high cases across all cancer types and for each cancer type.

### ChIP-Atlas analysis

Genome-wide transcription factor binding site information was obtained from chromatin immunoprecipitation (ChIP)-Atlas, an integrative database that covers public ChIP-seq data submitted to the NCBI SRA^34^. Under PeakBrowser, IRF3 protein binding was visualized at a given genome locus TSS using IGV genome browser^28^ using reference murine (version mm10) and human (version: hg38) genome.

### Statistical analyses

Data are presented as means ± standard error of the mean (SEM) from at least three independent experiments. For TCGA RNA-sequencing based gene expression analyses, comparisons of immune-related scores between *SMARCA4*-low vs *SMARCA4*-high expressing cancers were performed using the Mann-Whitney U test, followed by p-value adjustment using Benjamini-Hochberg method for multiple comparisons^23^ where p<0.05 was considered significant. For qRT-PCR and flow cytometry data analyses, comparisons between two groups (sg*SMARCA4* vs sgNTC) were determined by unpaired t-test comparison test. P-values <0.05 were considered significant. For RNA-sequencing data analysis for non-coding genes, statistical significance was considered if the adjusted P-value (after Benjamini-Hockberg correction) was p<0.05. For tumor growth data, studies were analyzed by first reducing the entire data on a given animal to a single number, the area under the curve, and then comparing the groups using the AUCs as data points, via a Wilcoxon test. When two independent experiments were pooled, the statistical test was stratified by the experiment. P-values <0.05 were considered significant. Statistical analyses were done using GraphPad Prism version 9 (GraphPad Software Inc., http://graphpad.com). Flow cytometry data were analyzed using FlowJo software (Tree Star Inc.).

## Results

### Pan-cancer association of SMARCA4 expression with immune expression signatures

To investigate whether levels of *SMARCA4* gene expression are associated with differential immune signatures in cancer, we compared immune-related properties defined by Thorsson et al.^33^ between *SMARCA4*-low and *SMARCA4*-high expressing cancers across 29 tumor types (PanCancerAtlas dataset^32^). *SMARCA4*-low (lowest quartile) and *SMARCA4*-high (highest quartile) cases were defined within each tumor type to avoid confounding factors (see Methods).

Comparison between *SMARCA4*-low (n=∼1900) and *SMARCA4*-high cases (n=∼1900) expressing tumors across cancer types revealed a statistically significant increase in key immune scores, including IFN gamma response (p<0.0001; q<0.0001) and lymphocyte infiltration signature (p<0.0001; q<0.0001) scores, in *SMARCA4*-low tumors (**Fig 1A**). Leukocyte fraction (p<0.0001; q<0.0001) as well as pro-inflammatory immune cell types, such as M1-macrophages (p=0.022; q=0.044), Th1 cells (p<0.0001; q<0.0001) and Th17 cells, (p=0.0002; q=0.0007), were statistically significantly more prominent in the *SMARCA4*-low compared to the *SMARCA4*-high expressing cancers (**Fig 1B**). By contrast, immune cell populations associated with inhibition of anti-tumor immunity such as Th2 cells (p<0.0001; q<0.0001) and regulatory T cells (p<0.0001; q<0.0001) were decreased in the *SMARCA4*-low compared to *SMARCA4*-high expressing cancers (**Fig 1C**).

**Figure 1.**
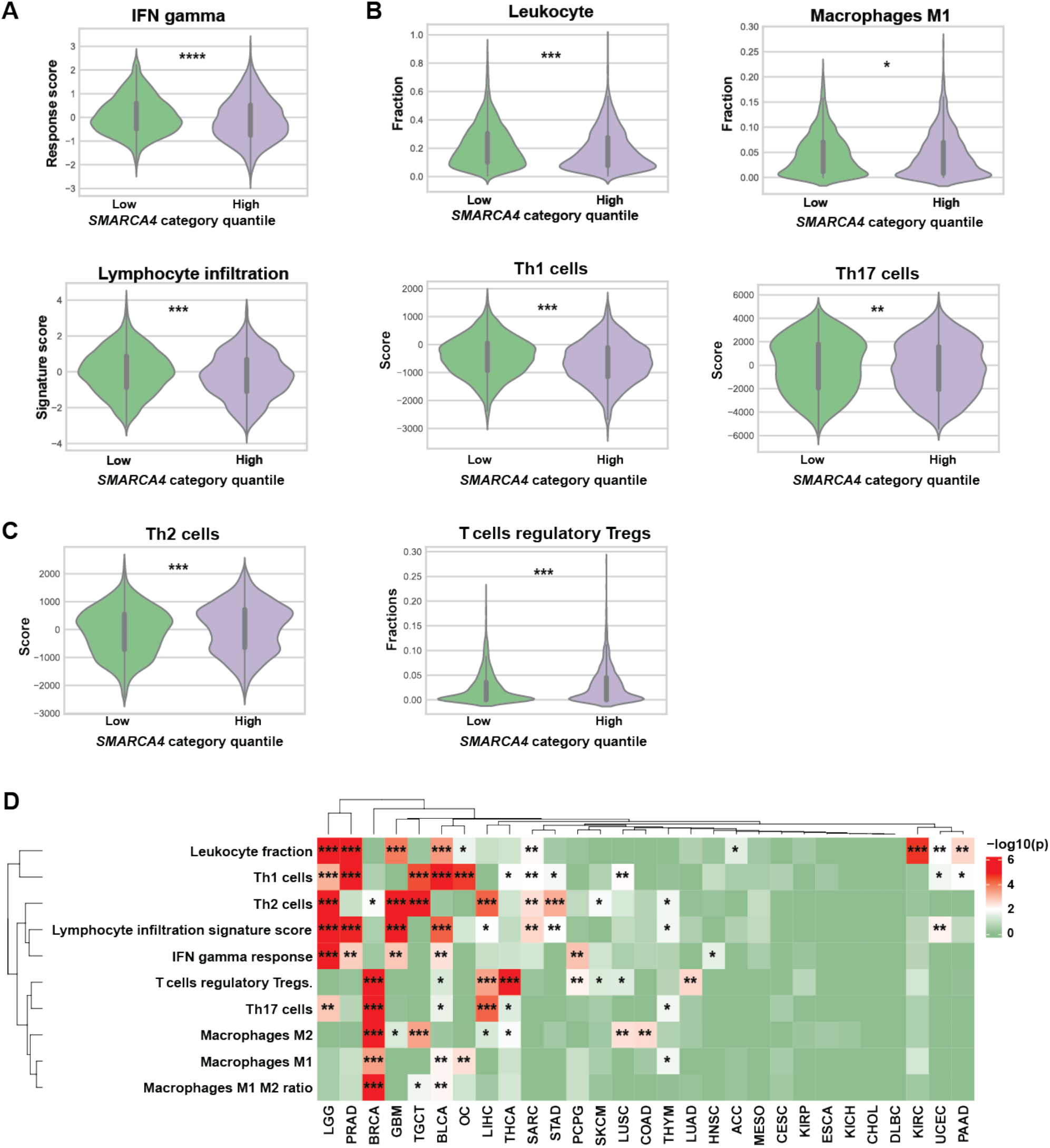
Pan-cancer association of *SMARCA4* expression with immune expression signatures. Transcriptomic analysis of The Cancer Genome Atlas (TCGA) pan-cancer RNA-sequencing data. **A)** Immune signatures (from Thorsson et al.^33^ dataset) by *SMARCA4* expression for all tumor types combined. **B)** Signature scores for pro-inflammatory immune cell populations by *SMARCA4* expression for all tumor types combined. **C)** Signature scores for immunomodulatory cell populations by *SMARCA4* expression for all tumor types combined. **D)** Heatmap of immune signatures in *SMARCA4*-low and *SMARCA4*-high tumors for each tumor type. Gene expression fold-change of *SMARCA4*-low versus *SMARCA4*-high tumors is color-coded according to the legend. Statistical analysis was performed using Mann-Whitney U test and Benjamini-Hochberg adjustment. *P<0.05, **P<0.01, ***P<0.001, ****P<0.0001. ACC=adrenocortical carcinoma, BLCA=bladder urothelial cancer, BRCA=breast invasive carcinoma, CESC=cervical squamous cell carcinoma and endocervical adenocarcinoma, CHOL=cholangiocarcinoma, COAD=colon adenocarcinoma, DLBC=lymphoid neoplasm diffuse large B-cell lymphoma, ESCA=esophageal carcinoma, GBM=glioblastoma multiforme, HNSC=head and neck squamous cell carcinoma, KIRC= kidney renal clear cell carcinoma, KIRP= kidney renal papillary cell carcinoma, LGG= brain low grade glioma, LIHC=liver hepatocellular carcinoma, LUAD=lung adenocarcinoma, LUSC= lung squamous cell carcinoma, MESO=mesothelioma, OC=Ovarian serous cystadenocarcinoma, PAAD=pancreatic adenocarcinoma, PRAD=prostate adenocarcinoma, SARC=sarcoma, SKCM=skin cutaneous melanoma, STAD= stomach adenocarcinoma, TGCT= testicular germ cell tumors, THYM=thymoma, UCEC=uterine corpus endometrial carcinoma.

Similar analyses by cancer type demonstrated associations of immune signatures and immune cell infiltration with *SMARCA4* expression across a number of tumor types (**Fig 1D**). In OC (n=171), higher IFN gamma response and lymphocyte infiltration signature score was similarly observed in the *SMARCA4-*low (n=43) cohort. Leukocyte fraction (p=0.003; q=0.024) was increased in *SMARCA4*-low OCs with prominence of M1-macrophages (p=0.0004; q=0.003) and Th1 cells (p<0.0001; q<0.0001) (**Fig 1D, Fig S2**).

We also explored the prevalence of various type of *SMARCA4* mutations in our predefined *SMARCA4*-low and *SMARCA4*-high groups across cancer types. Truncating and splice-site mutations were significantly more frequent in the *SMARCA4*-low than *SMARCA4*-high expressing cancers (p<0.0001; **Table S7**). In contrast, cancers with missense *SMARCA4* somatic mutations were similarly distributed between *SMARCA4*-low than *SMARCA4*-high expressing cancers (p=0.0527; **Table S7**). Overall, these data support the rationale for studying the impact of loss-of-function mutations in *SMARCA4* on the immune response.

### Loss of function of SMARCA4 in OC cells leads to increased interferon response and antigen presentation gene activation

We generated *SMARCA4*-knockout (sg*SMARCA4*) murine (ID8 and UPK10) and human (OAW28) stable OC cell lines as well as stable non-targeted control for each (sgNTC). RNA-sequencing of the generated cell lines was employed to examine differentially expressed coding genes (**Fig S3A**) in sg*SMARCA4* and sgNTC OC cells. Hallmark and Gene Ontology enrichment analyses identified significant upregulation of IFN-alpha, -beta and -gamma signaling pathways as well as genes related to NFkB signaling and antigen processing/presentation in sg*SMARCA4* as compared to sgNTC cells (**Fig 2A, Fig S3B**). When specifically focusing on type I IFN pathway-related genes, we observed significant increase in basal levels of interferon-stimulated genes (ISGs) including *OASL2, ISG15, IRF7, IFI44L, IFI27, MX1* and *IFI44* in sg*SMARCA4* cells (**Fig 2B, Fig S3C**). By contrast, type III interferon genes were not upregulated in sg*SMARCA4* cells (**Fig S3D**). These results were validated by qRT-PCR (**Fig 2C, Fig S3E**).

**Figure 2.**
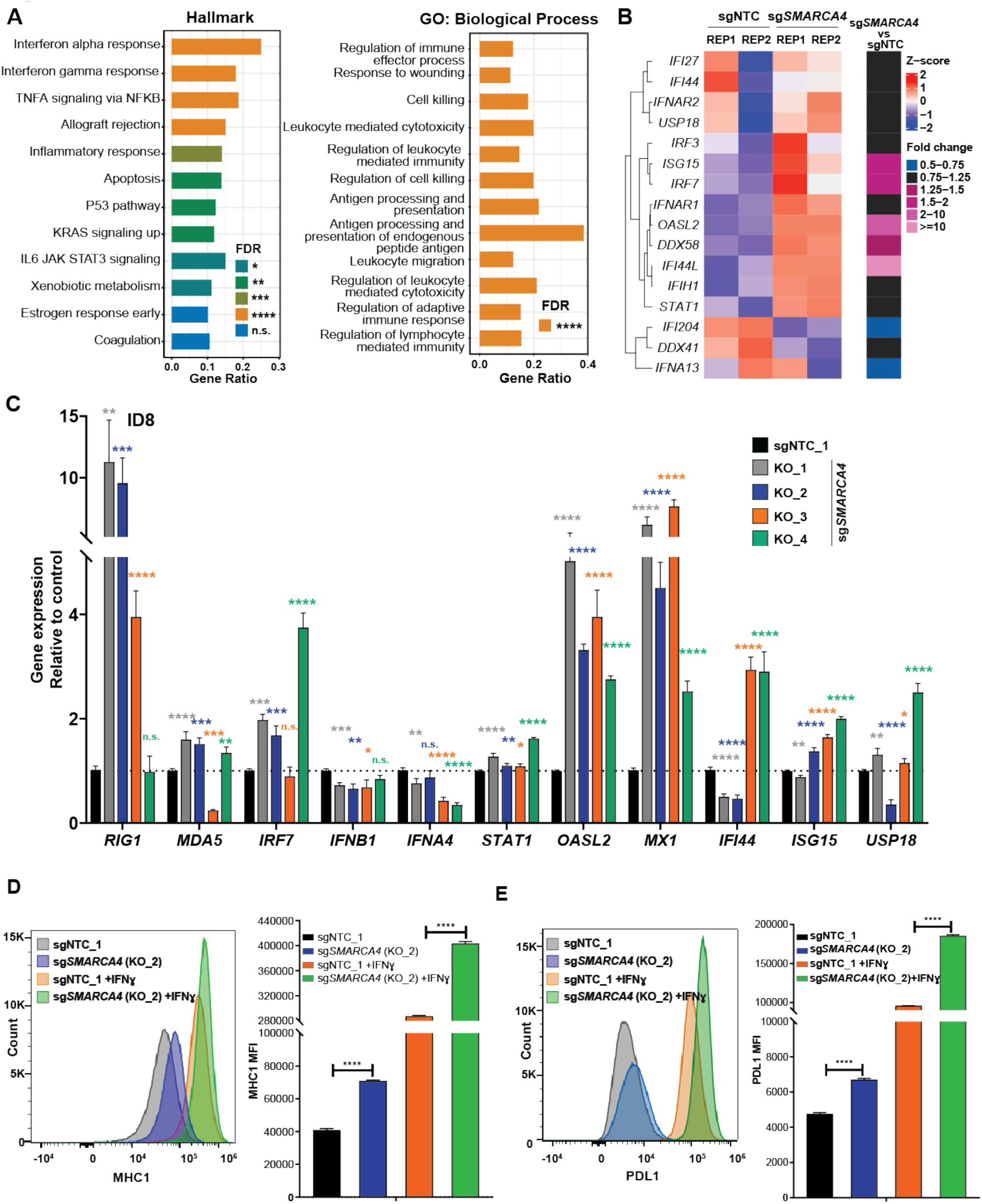
Loss of function of *SMARCA4* in ovarian cancer cells leads to increased interferon response and antigen presentation gene activation. **A)** Upregulated Hallmark and Gene Ontology pathways in ID8 cells single guide *SMARCA4* (sg*SMARCA4*) compared to sgNTC. Ribodeplete RNA-sequencing was performed. Statistical analysis was based on hypergeometric test and performed using ClusterProfiler. **B)** Gene expression heatmap of type I IFN pathway-related genes in ID8 cells. Reads Per Kilobase of transcript per Million mapped reads (RPKM) values were scaled to Z-score for visualization. Gene expression fold-change of sg*SMARCA4* versus sgNTC cells is color-coded according to the legend. **C)** qRT-PCR validation results for IFN genes in ID8 cells (sgNTC + 4 sg*SMARCA4* clones). Expression levels were normalized to B-actin expression, and comparisons of mRNA expression levels were performed relative to control (sgNTC). N=3 independent experiments. **D)** MHC1 expression in ID8 cells with or without IFNɣ by flow cytometry. **E)** PDL1 expression in ID8 cells with or without IFNɣ by flow cytometry. Statistical analysis was performed using two-tailored unpaired t-test (**C, D, E**). *P<0.05, **P<0.01, ***P<0.001, ****P<0.0001, n.s. = not significant. Error bars represent ± SEM. Samples in duplicates (for **A, B**) and triplicates (for **D, E**). GO = Gene Ontology, IFN = interferon, KO = knockout, NTC = non-target control.

We next examined the effects of *SMARCA4* loss on the downstream immune signaling, including antigen presentation and immune checkpoint expression. MHC1 expression assessed by flow cytometry was significantly increased in sg*SMARCA4* compared to sgNTC OC cells, both at baseline and in the setting of IFNɣ stimulation (**Fig 2D, Fig S3F**). Similarly, at baseline sg*SMARCA4* cells expressed higher levels of PD-L1 than sgNTC cells and exhibited further upregulation of PD-L1 in the presence of IFNɣ stimulation (**Fig 2E**). Overall, these results highlight that loss of *SMARCA4* leads to alteration of tumor cell-intrinsic immune signaling, potentially impacting both innate (type I IFN) and downstream adaptive (antigen presentation) immune responses.

### SMARCA4 loss of function mutation results in perturbation of chromatin accessibility at key immune genes

Given that *SMARCA4* is a core member of the mSWI/SNF chromatin remodeling complex, we hypothesized that *SMARCA4* loss alters chromatin accessibility for IFN-related gene transcription. We used the ATAC sequencing (ATAC-Seq) to assess and compare chromatin accessibility changes in sg*SMARCA4* or sgNTC OC cells. Loss of *SMARCA4* led to decrease in accessibility in some sites and increase in others (**Fig 3A**). Immune and chemokine related pathways were found to be enriched in genes with increased accessibility (**Fig 3B**). Furthermore, we found significant increase of chromatin accessibility at Th1-type chemokine loci in sg*SMARCA4* cells, such as CXCL10 (*q*= 0.02) and CCL2 (*q*<0.0001) genes, with strong peaks on their transcription start sites (TSS) (**Fig 3C**). This was consistent with upregulation seen in RNA-sequencing data. We further performed motif enrichment analysis, and identified that the AP1, NFκB and IRF motifs, which are bound by transcription factors that regulate innate and IFN-stimulated genes, are enriched in the open chromosomal regions affected by *SMARCA4* deficiency (**Fig 3D**). Overall, these results suggest that loss of *SMARCA4* leads to changes in chromatin accessibility associated with induction of IFN-response genes.

**Figure 3.**
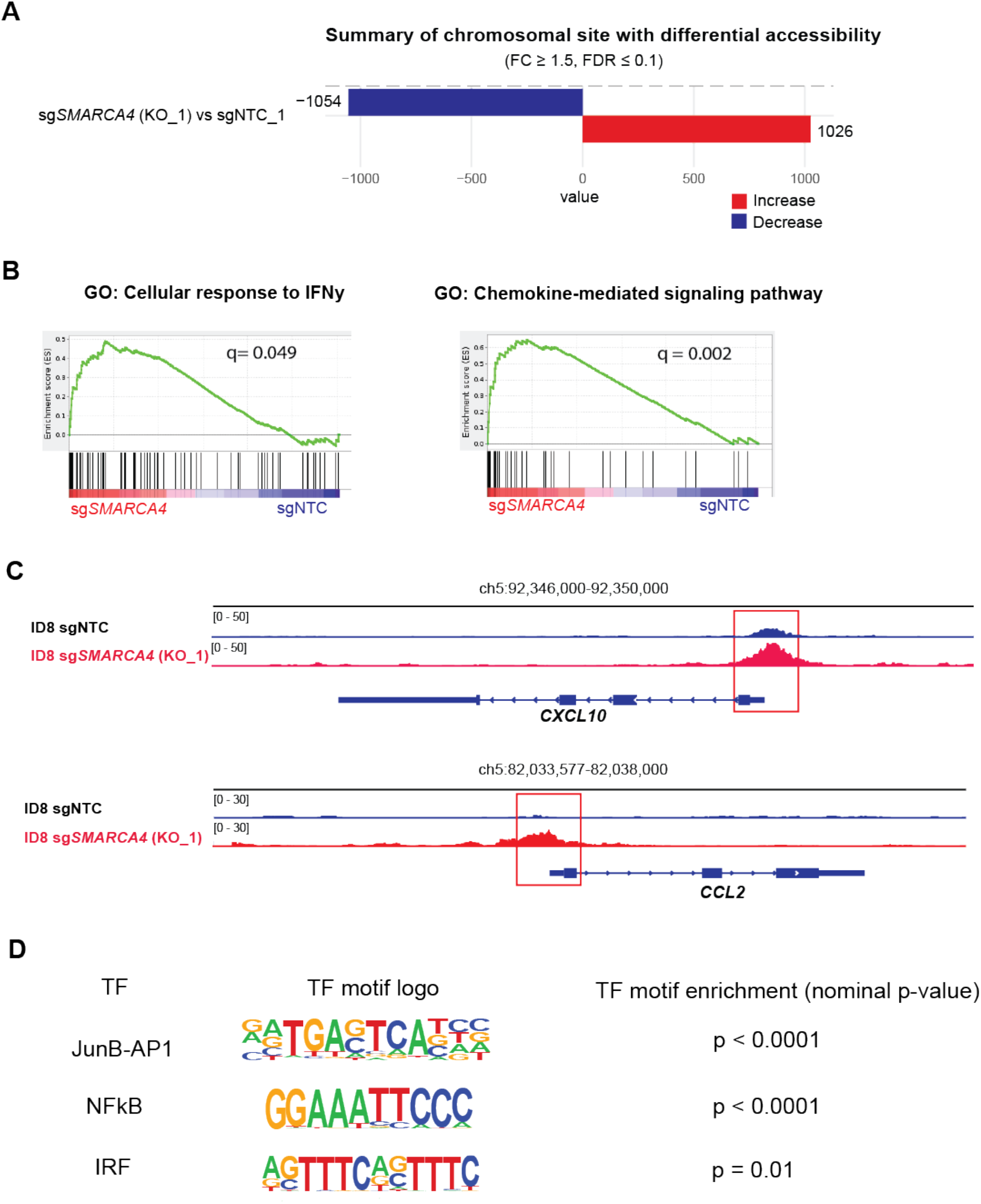
*SMARCA4* loss of function mutation results in perturbation of chromatin accessibility at key immune genes. **A)** Changes in genomic site accessibility in sg*SMARCA4* versus sgNTC cells. Log2 fold change and FDR-adjust p-value (Wald test p-values from DESeq2 with Benjamini-Hochberg correction). **B)** Gene set enrichment analysis of immune pathways with increased accessibility in sg*SMARCA4* versus sgNTC cells. Kolmogorov-Smirnov statistic with Benjamini-Hochberg correction. Exact q values indicated in each panel. **C)** Changes in chromatin accessibility at transcription start sites of ISGs (CXCL10, CCL2) in sg*SMARCA4* versus sgNTC cells. **D)** Motifs enriched in open chromosomal regions affected by *SMARCA4* deficiency. P-value and binomial test were performed using Homer2. Experiments performed in duplicates. GO = Gene Ontology, ISG = interferon stimulated gene, NTC = non-target control.

### Increase in ISGs is not dependent on signaling through the type I IFN receptor in **sg*SMARCA4 OC cells***

We initially hypothesized that upregulation of ISGs was mediated through enhanced signaling through type I IFN receptor (IFNAR). While IFNAR blockade with anti-IFNAR antibody resulted in attenuation of ISG expression in response to type I IFN stimulation (**Fig S4A**), surprisingly, blocking the IFNAR did not attenuate the ISG expression in ID8 sg*SMARCA4* cells (**Fig 4A**). Interestingly, in the human ovarian sg*SMARCA4* OAW28 cell line, blocking IFNAR resulted in partial attenuation of ISG expression (**Fig S4B**). These findings suggest that in sg*SMARCA4* cells transcription activation of ISGs is not fully dependent on activation of the IFNAR by type I IFN, though the discrepancy between the murine and human cells indicate that this dependency may be cell line-or species-specific.

**Figure 4.**
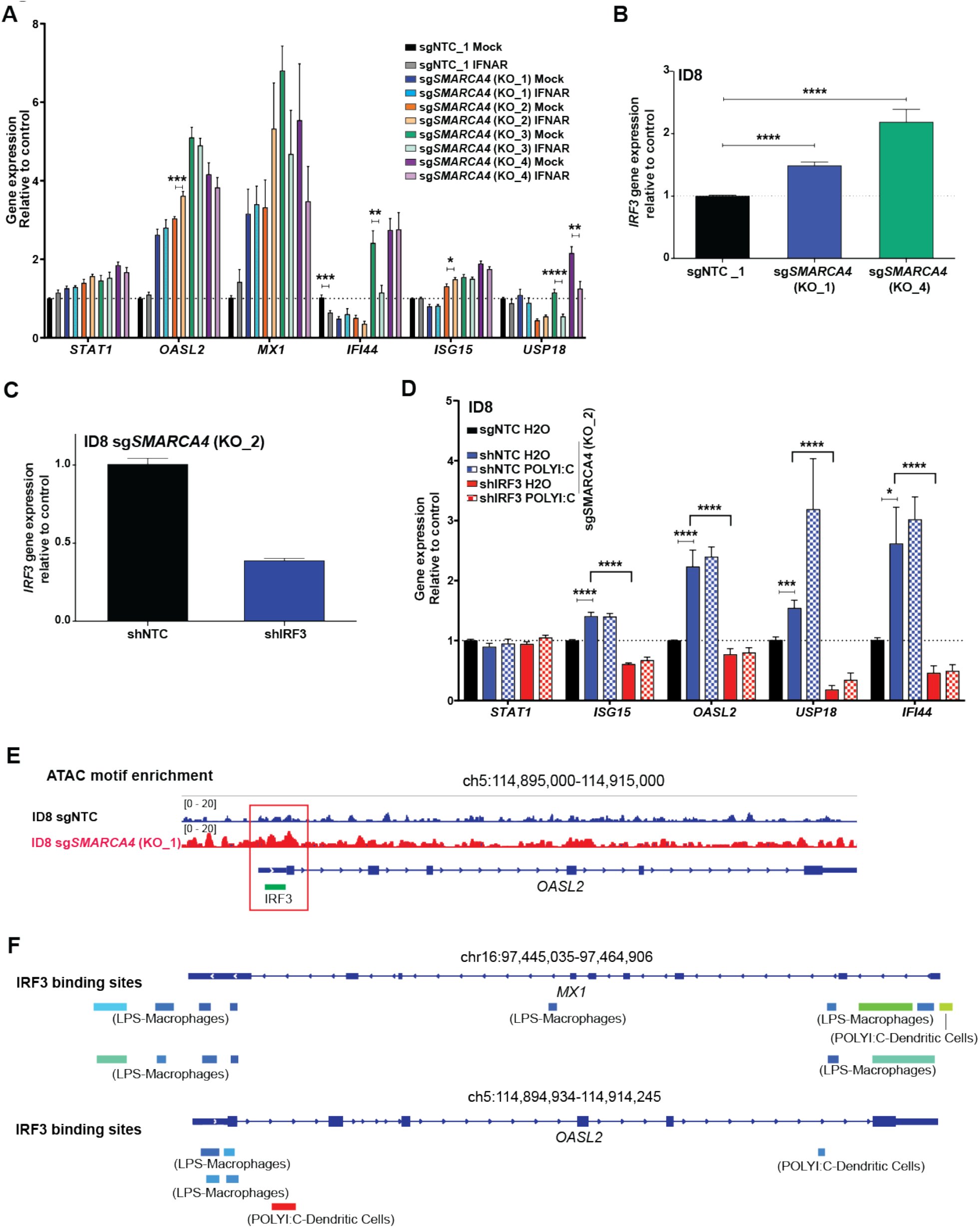
Increase in ISGs is not dependent on signaling through the type I IFN receptor in sg*SMARCA4* OC cells. **A)** IFNAR-1 neutralizing assay in ID8 single guide NTC (sgNTC) compared to sg*SMARCA4* cells (4 sg*SMARCA4* clones). Cells were treated with Mock or IFNAR-1 antibody for 48 hours at 10 ug/mL. **B)** IRF3 expression levels in sg*SMARCA4* vs sgNTC cells by qRT-PCR. **C)** IRF3 expression in doxycycline-inducible short hairpin NTC (shNTC) and shIRF3 transfected sg*SMARCA4* cells by qRT-PCR. **D)** qRT-PCR quantification of ISGs in ID8 sgNTC cells and ID8 sg*SMARCA4* cells. The latter were transfected with shIRF3 or shNTC (with and without POLYI:C). **E)** Enriched IRF motif at transcription start sites of ISG locus from ATAC-seq analysis of ID8 sgNTC versus sg*SMARCA4* cells. Experiments performed in duplicates. **F)** Analysis of publicly available ChIP-Atlas data of IRF3 DNA binding sites on ISGs in murine immune cell lines. Statistical analysis was performed using two-tailored unpaired t-test (**A, B, C, D**). *P<0.05, **P<0.01, ***P<0.001, ****P<0.0001. Error bars represent ± SEM. N=3 independent experiments in **A, B, C, D**. Expression levels were normalized to B-actin expression, and comparisons of mRNA expression levels were performed relative to control (sgNTC in **A, B, D** and sg*SMARCA4* in **C**, as indicated). ISG = interferon stimulated gene, LPS = lipopolysaccharide stimulated, NTC = non-target control, POLYI:C = Polyinosinic-polycytidylic acid stimulated.

We proceeded to evaluate whether activation of ISGs is dependent on the Interferon Regulatory Factor 3 (IRF3), a key upstream regulator of type I IFN gene transcription. Loss of *SMARCA4* was associated with increase in IRF3 RNA and protein expression (**Fig 4B, Fig S4C**) and nuclear translocation (**Fig S4D**). Knocking down IRF3 (**Fig 4C, Fig S4E**) abrogated ISG upregulation in sg*SMARCA4* cells (**Fig 4D**) confirming that ISG activation is at least partially dependent on IRF3.

We examined the IRF3 binding sites at the ISG promoters as a potential mechanism that could explain the IFNAR-independent ISG upregulation that we observed with *SMARCA4* loss. Using the ATAC-seq data, we found that IRF3 motifs are enriched at the TSS of ISG locus (*OASL2*), suggesting that IRF3 may bind at a response element locus at the promoter site of ISGs (**Fig 4E**). Analysis of publicly available datasets from ChIP-Atlas revealed IRF3 binding sites at the promoters of key ISGs including *MX1* and *OASL2* in human cancer cell lines (**Fig 4F**) as well as key ISGs such as *OASL, MX1* and *MX2* in murine immune cell lines with or without immunostimulants (**Fig S4F**). Put together, these results suggest that in the setting of *SMARCA4* deficiency, upregulation of ISGs is at least in part driven by IRF3, potentially through increased IRF3 expression or upstream activation.

### Increase in ISGs is mediated through the dsRNA sensing pathway

Next, we focused on the potential upstream signaling mechanisms leading to IRF3 activation. We hypothesized that chromatin remodeling associated with loss of *SMARCA4* could lead to re-expression of noncoding and transposable elements (TE) and tandem repeats which carry a potential to engage innate receptors that have been shown to lead to type I IFN pathway activation in other settings^7,11,35^. To address this, we assessed the differentially expressed TEs in the transcriptomes of sg*SMARCA4* versus sgNTC cells. Loss of *SMARCA4* in the murine ID8 cells induced a significant increase in long terminal repeats (LTRs) type of TEs which include endogenous retroviruses (ERVs) (**Fig 5A, B**), which was confirmed in the UPK10 model using qRT-PCR of a panel of ERV genes (**Fig S5A**). This ERV increase was not observed in the human OAW28 sg*SMARCA4* cells (**Fig S5B, C**). In contrast, long and short interspersed nuclear elements (LINEs, SINEs) TEs were generally downregulated in the sg*SMARCA4* cells compared to sgNTC cells (**Fig 5A, Fig S5B**). Tandem satellite repeats (SATs) were upregulated in OAW28 sg*SMARCA4* cells compared to sgNTC cells (**Fig S5D**) but remained unchanged in ID8 sg*SMARCA4* cells (**Fig 5A**).

**Figure 5.**
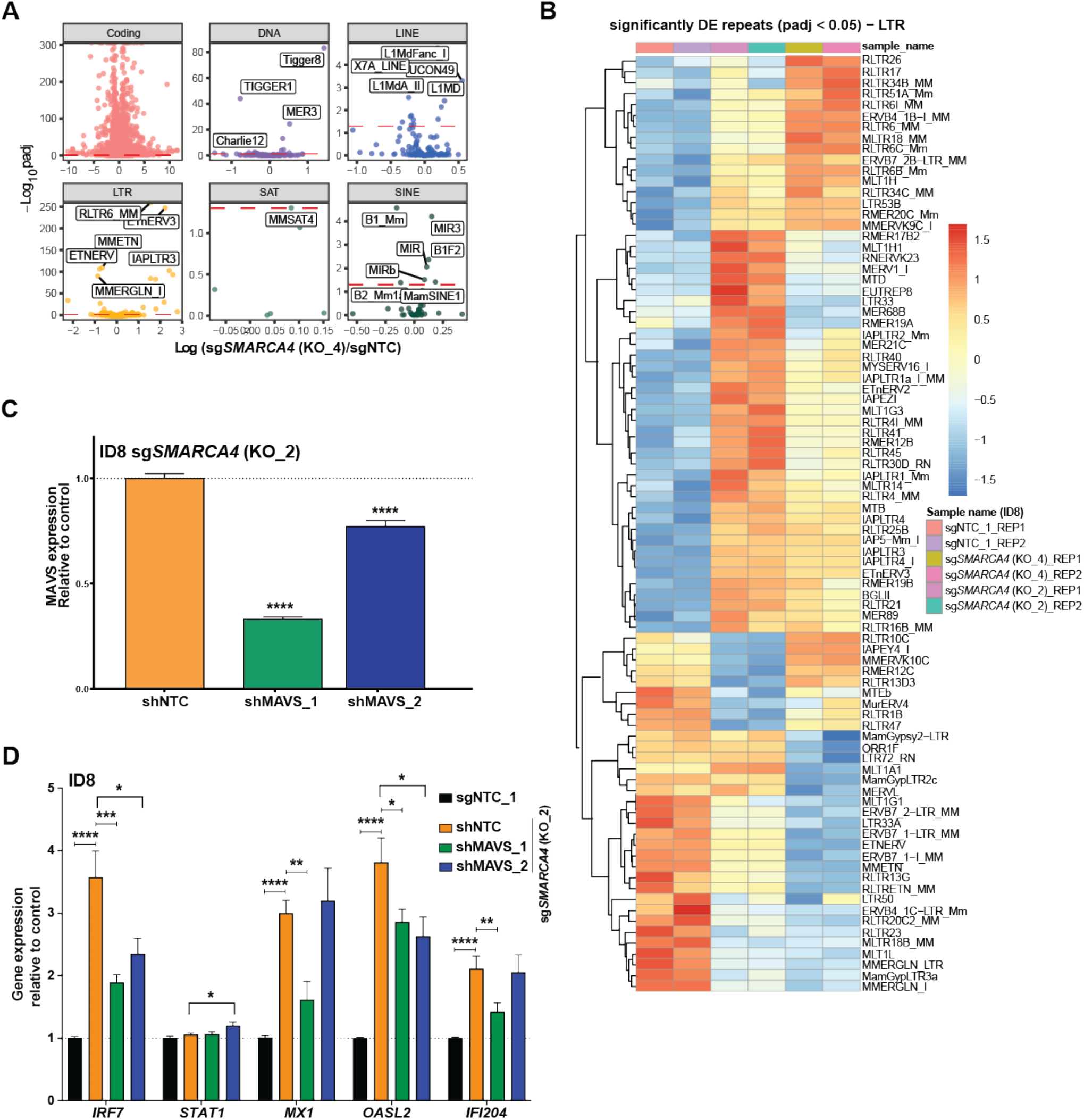
Increase in ISGs is mediated through the dsRNA sensing pathway. **A)** Volcano plots for differential expression of TEs in ID8 cells. **B)** Heatmap of long terminal repeats (LTR) repeats with adjusted p<0.05. **C)** MAVS expression in doxycycline-inducible shNTC and shMAVS transfected sg*SMARCA4* cells by qRT-PCR. **D)** qRT-PCR quantification of ISGs in ID8 sgNTC cells and ID8 sg*SMARCA4* cells transfected with shMAVS or shNTC. Expression levels were normalized to B-actin expression, and comparisons of mRNA expression levels were performed relative to control (sg*SMARCA4* in **C** and sgNTC in **D**, as indicated). Statistical analysis was performed using two-tailored unpaired t-test (**C, D**). *P<0.05, **P<0.01, ***P<0.001, ****P<0.0001. Error bars represent ± SEM. Samples in duplicates (for **A, B**). N=3 independent experiments (in **C, D**). Red line in (**A**) represents a cut-off of an adjusted p-value <0.05. ISG = interferon stimulated gene, NTC = non-target control.

We proceeded to determine whether the RNA sensing RIG-I-like receptor (RLR) pathway could be potentially driving ISG upregulation, possibly through detection of increased LTR expression in sg*SMARCA4* cells. We performed shRNA knockdown of the mitochondrial antiviral signaling (MAVS) protein, a key downstream adapter mediating RIG-I and MDA5 signalling (**Fig 5C, Fig S5E**). Knockdown of MAVS resulted in downregulation of expression of different ISGs in sg*SMARCA4* ID8 cells (**Fig 5D**). Taken together, these data suggest that ISG upregulation induced by loss of function of *SMARCA4* is at least in part triggered by the RNA sensing pathway, possibly through increased expression of repeat elements as has been previously demonstrated within the setting of hypomethylating agent therapy^7^.

### sgSMARCA4 leads to increase of tumor immunogenicity in vivo

We proceeded to examine the impact of *SMARCA4* loss on the immune response *in vivo* in the peritoneal ID8 OC model (**Fig 6A, Fig S6, Fig S7**). While there was no significant change in the numbers of CD4+ and CD8+ T cells in the ascites of sg*SMARCA4* OCs (**Fig S8A**), we observed evidence of enhanced CD4+ and CD8+ T cell activation, as demonstrated by increased expression of PD1 (p<0.0001; **Fig 6B**) and increased but not statistically significant Granzyme B expression (**Fig S8B**). There was no substantial change in regulatory T cells between the sg*SMARCA4* and sgNTC ascites (**Fig S8C**). We also observed increased activation of NK cells as evidenced by Granzyme B expression (p=0.03; **Fig 6C**) though there was a relative decrease in the overall NK cell numbers (**Fig S8D**). Evidence of increased immunogenicity of *sgSMARCA4* cells *in vivo* was also observed in the B16-F10 melanoma model (**Fig S8E**), demonstrating increased CD3+ and CD4+ T cells (**Fig S8F**) as well as increased NK cell infiltration (**Fig S8G**).

**Figure 6.**
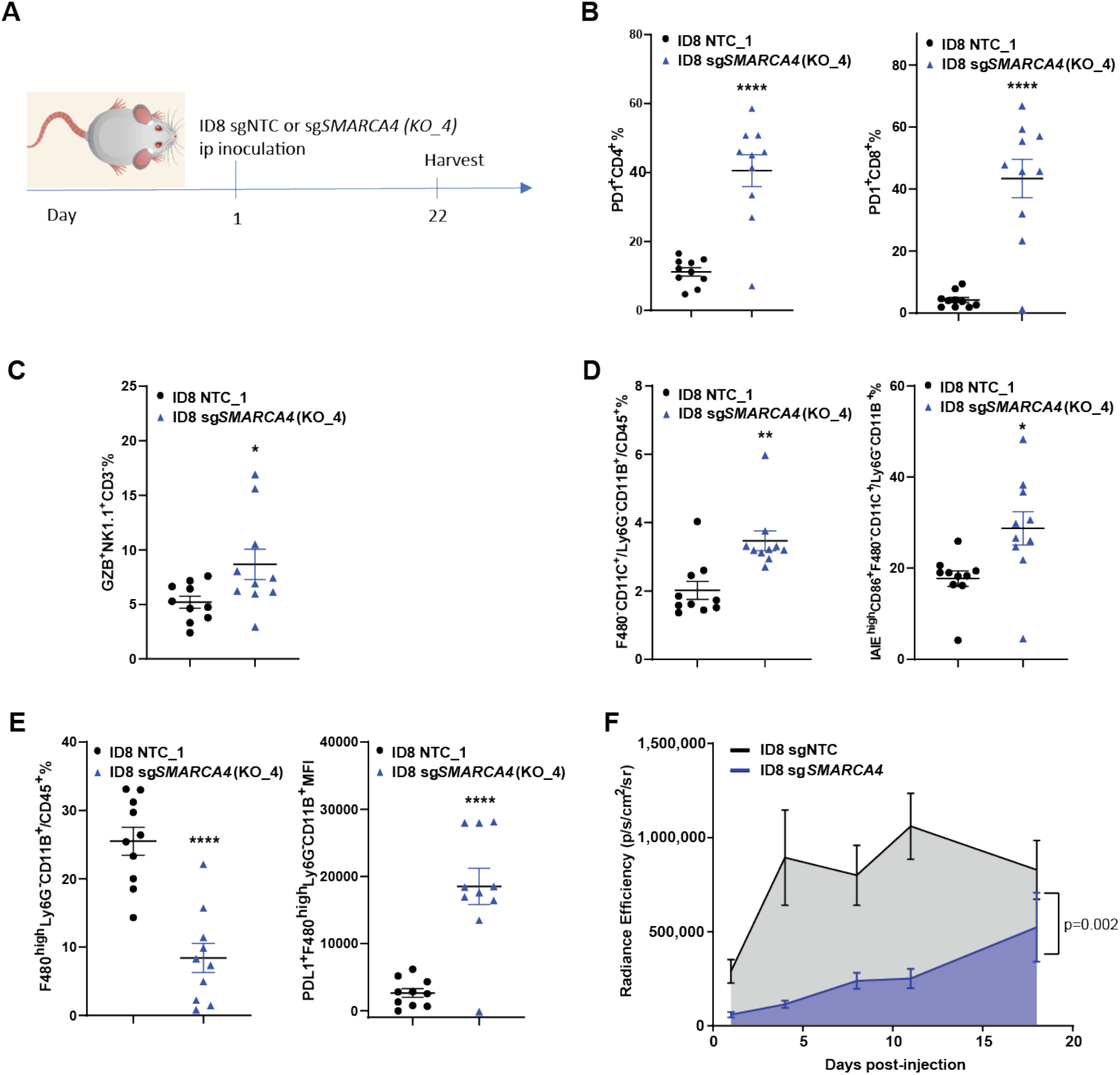
*SMARCA4* loss of function leads to increase immunogenicity *in vivo*. **A)** Workflow for the ovarian cancer tumor model. C57BL/6 (Cg)-Tyr^c-2J^/J mice were inoculated intraperitoneally with 10 million ID8 sgNTC or sg*SMARCA4* tumor cells and spectral flow cytometry was performed 21 days later. **B)** Frequency of tumor PD1+CD4+ T cells and PD1+CD8+ T cells. **C)** Frequency of NK1.1+ cells expressing Granzyme B+. **D)** Frequency of tumor dendritic cells (of CD45+ cells) and MHCII expressing dendritic cells. **E)** Frequency of tumor macrophages (of CD45+ cells) and PDL1 expressing macrophages. **F)** *In vivo* bioluminescence imaging of tumor burden in sg*SMARCA4* vs sgNTC tumors (unpaired t-test of area under the curve). Statistical analysis was performed using two-tailored unpaired t-test (**B, C, D, E, F**). *P<0.05, **P<0.01, ***P<0.001, ****P<0.0001. Error bars represent ± SEM. N=10 mice/group. IP= intraperitoneal, NTC = non-target control.

In the myeloid cell compartment, we observed significantly increased numbers and MHCII activation of dendritic cells (p=0.01; **Fig 6D**). Furthermore, there was a significant decrease in the overall number of macrophages (p<0.0001), though the remaining macrophages exhibited increased expression of PDL1 (**Fig 6E**). Similar findings were found in the B16-F10 model (**Fig S8H**). Finally, while the sg*SMARCA4* ID8 cell line growth *in vitro* was comparable to the sgNTC cells (**Fig S8I**), there was a substantial delay in tumor growth *in vivo* as assessed by bioluminescence imaging (p=0.002; **Fig 6F**). This was also validated in the B16-F10 model showing suppressed tumor growth in sg*SMARCA4* tumors (**Fig S8J**). Overall, we observe that loss of *SMARCA4* is associated with increased immunogenicity *in vivo*, as evidenced by a more active innate and adaptive immune TME and tumor growth delay.

### Therapeutic targeting of BRG1 recapitulates the immunogenic effects of sgSMARCA4

Our findings suggest that genetic loss of *SMARCA4* is associated with increased tumor immunogenicity, which led us to hypothesize that targeting *SMARCA4* therapeutically may also promote tumor immunogenicity. To mimic the loss of *SMARCA4*, a dual BRG1/BRM inhibitor (HY- 119374) and BRG1/BRM proteolysis-targeting chimera (PRTD-0043) were tested in *SMARCA4-* wildtype murine OC cell lines (ID8, UPK10). Treatment of cells with either compound led to upregulation of key type I IFN-related genes (**Fig 7A and Fig S9A, B**). We proceeded to assess treatment with the HY-119374 compound in ID8 tumors *in vivo* (**Fig 7B**). Treatment with the inhibitor resulted in increased CD3+ cells with significant increase in CD8+ T cells (p=0.02; **Fig 7C**) and a similar trend for CD4+ T cells (**Fig S9C**). Both CD4+ and CD8+ T cells exhibited higher levels of activation, as evidenced by upregulation of ICOS, PD1 and Granzyme B expression (**Fig 7D, E**). We similarly observed an increase in NK cell activation (**Fig 7F**) though the relative NK cell numbers remained unchanged (**Fig S9D**).

**Figure 7.**
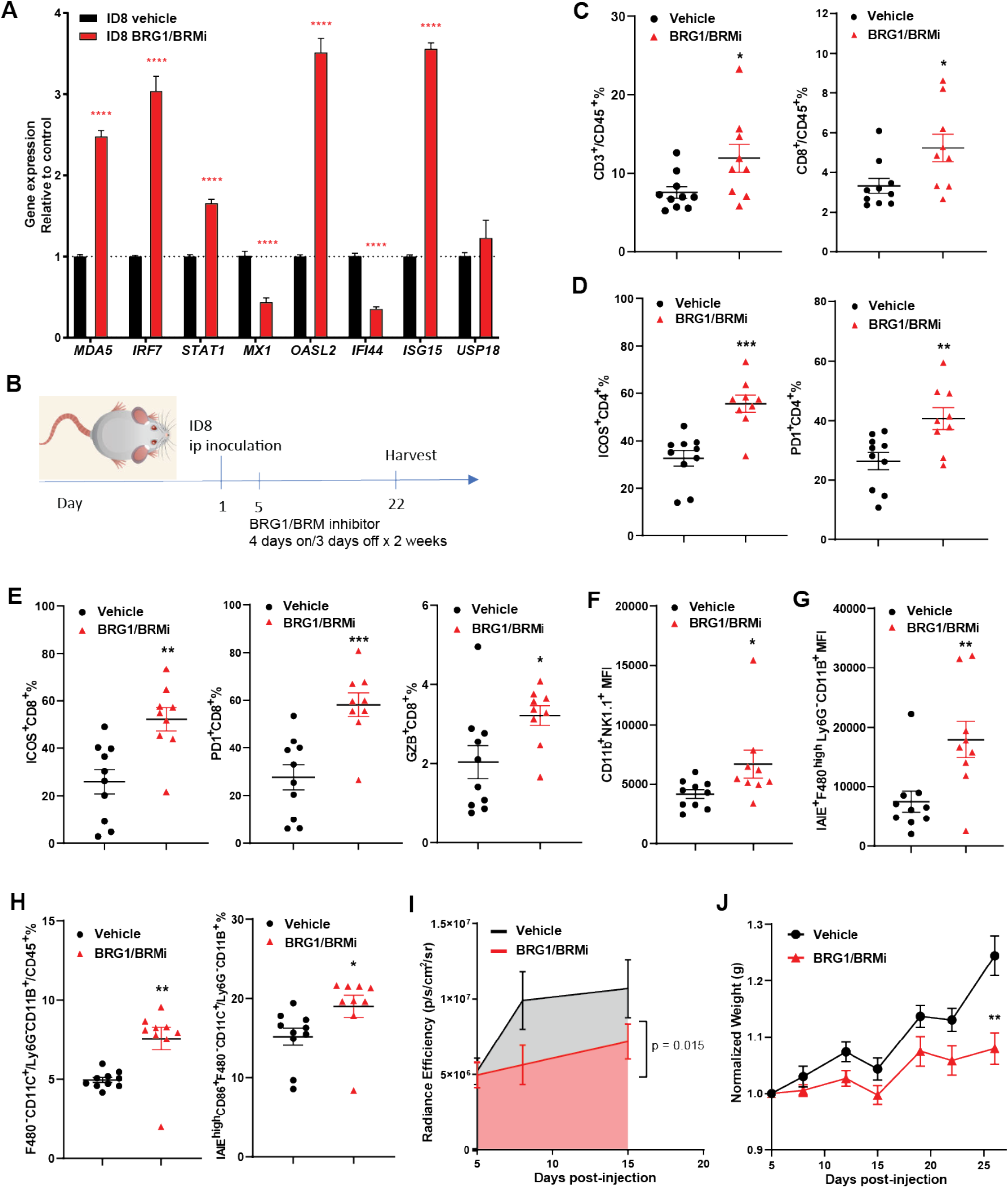
Therapeutic targeting of SMARCA4 recapitulates the immunogenic effects of *SMARCA4* loss. **A)** qRT-PCR quantification of IFN gene expression using ID8 parental cell treated with 1uM of BRG1/BRM inhibitor or vehicle (DMSO). Expression levels were normalized to B-actin expression, and comparisons of mRNA expression levels were performed relative to control (vehicle). **B)** Workflow for the ovarian cancer tumor model and treatment. C57BL/6 mice were inoculated IP with 2 million ID8 parental cells and treated with 20 mg/kg of BRG1/BRM inhibitor or vehicle Solutol HS (10 mM citrate buffer, pH6 in a ratio of 5:20:75) starting on Day 5 for a total of 8 doses (4 days on, 3 days off x 2 weeks). Flow cytometry was performed 3 weeks post-inoculation. **C)** Frequency of tumor CD3+ cells and CD8+ T cells (of CD45+ cells). **D)** Frequency of tumor ICOS+CD4+ T cells and PD1+CD4+ T cells. **E)** Frequency of tumor ICOS+CD8+ T cells, PD1+CD8+ T cells and Granzyme B expressing CD8+ T cells. **F)** MFI of activated (CD11b+) NK1.1+ cells. **G)** Frequency of tumor dendritic cells (of CD45+ cells) and MHCII expressing dendritic cells. **H)** MFI of MHCII expressing tumor macrophages. **I)** *In vivo* bioluminescence imaging of tumor burden in vehicle vs BRG1/BRMi-treated tumors (unpaired t-test of area under the curve). **J)** Mice weights in ID8 tumors treated with BRG1/BRMi or vehicle. Weights normalized to baseline weight (Day 5). Statistical analysis was performed using two-tailored unpaired t-test. *P<0.05, **P<0.01, ***P<0.001, ****P<0.0001. Error bars represent ± SEM. N=10 mice/group in **C, D, E, F, G, H, J** and N= pooled 20 mice/group from two independent experiments in **I**. BRG1/BRMi = BRG1/BRM inhibitor, IP = intraperitoneal

In the myeloid compartment, there was no significant change in macrophages, however we observed a significant increase in macrophage MHCII expression (p=0.0077; **Fig S9E, Fig 7G**). A significant increase was noted in dendritic cells and upregulation of MHCII expression on their surface (**Fig 7H**). Myeloid-derived suppressor cells (defined by F4/80^intermediate/negative^Ly6C^-^CD206^+^ phenotype) were overall decreased (**Fig S9F**) reflecting a more pro-inflammatory TME. Lastly, treatment with HY-11934 significantly decreased tumor burden (p=0.015; **Fig 7I**) and inhibited ascites accumulation (**Fig 7J, Fig S9G**) further supporting the anti-tumor immune effect of loss of function of *SMARCA4*. Overall, these results demonstrate that therapeutic inhibition of BRG1 leads to immune activation in the TME, supporting the rationale for this therapeutic strategy as an approach to increase tumor immunogenicity and make them potentially more susceptible to T cell-targeted immunotherapies like ICB.

## Discussion

Traditionally, tumor immunogenicity and response to ICB have been correlated with biomarkers such as TMB and PDL1 expression^4^, neither of which carry strong predictive value in OC^36^. Several studies have suggested that disruption in chromatin remodelling may serve as a predictor of immunogenic properties of cancers^5,12,37^. Proficient chromatin remodelling has been associated with T cell exhaustion and poor tumor control^38^ and conversely, depletion of chromatin regulators correlated with increased ICB sensitization^9^. Previous preclinical murine data also suggest that SWI/SNF-deficient tumors, including loss-of-function of *SMARCB1*, *PBRM1* and *ARID1A*, respond better to ICB^11,12,39^. Clinical results also support these preclinical findings including in *SMARCA4*-deficient lung cancers^13^, *PBRM1*-mutated clear cell renal cell carcinoma^40^ and clear cell OCs^41^ of which over 50% have SWI/SNF mutations^15^.

Overall, the impact of epigenetic dysregulation in OC on the immune TME remains relatively uncharacterized. Our study results provide a mechanistic understanding of observations made from previous published work suggesting improved ICB response^5,13^ and immunogenic OC TME in *SMARCA4*-deficient cancers^5^. We demonstrate that loss of *SMARCA4* leads to activation of innate immune signaling leading to IRF3-dependent induction of downstream ISGs, chemokines and antigen presentation machinery. The type I IFN pathway is responsible for inducing transcription of a large group of genes which play a role in response to viral infection as well as activating key components of the innate and adaptive immune systems such as antigen presentation and production of immunostimulatory cytokines.

We find that this immune activation may be in part triggered by activation of the RLR pathway. We also show that *SMARCA4* loss leads to re-expression of normally silenced TEs such as LTRs. Though our study does not establish a direct link between TE expression and RLR activation, several studies in the past have demonstrated that de-repression of repeat elements within the context of hypomethylating agent therapy can drive activation of RLR-dependent activation of type I IFN pathway^7,42^. In addition, although traditionally categorized as non-coding proteins, recent studies have suggested that induced TEs may in fact be translated into proteins that can serve as neoantigens^9^.

Consistent with the increase in tumor cell-intrinsic immunogenicity in response to *SMARCA4* loss, we observe increased immune activation *in vivo*, as evidenced by delayed tumor growth, increase in activated T and NK cells, and increased infiltration of activated antigen presenting cells. Finally, we demonstrate the potential therapeutic application of these findings, by showing that a BRG1/BRM inhibitor mimicked the changes seen in sg*SMARCA4* tumors, resulting in increased infiltration with activated T and NK cells and antigen presenting cells.

Other epigenetic therapies have recently been shown to enhance tumor immunogenicity^7,43,44^. DNA methyltransferase inhibitors with or without histone deacetylase inhibitors have been shown to induce type I IFN genes leading to activation of innate and adaptive immune responses in the TME^7,43^. CARM1 (a protein that methylates and displaces BAF155, a subunit of the SWI/SNF complex) was found to be a negative regulator of tumor infiltrating T cells and, when ablated genetically or therapeutically, this led to increased tumor CD8+ T cell infiltration and activation as well as immune-related tumor suppression^44^. Further analyses showed that these effects were mediated by type I IFN induction and a heightened response to IFNα/γ^44^.

Overall, our findings demonstrate an immunomodulatory role of *SMARCA4* and generate rationale for further evaluation of *SMARCA4* loss of function mutations as predictors of response to immunotherapy and for use of SMARCA4/BRG1 as a therapeutic target to increase OC immunogenicity.

## Acknowledgements

We thank the staff of the research staff at Core Facilities at MSK: Marco Russo and Ralph Garippa (Gene Screening and Editing), Eric Rosiek and Eric Chan (Molecular Cytology), Cassidy Cobbs (Integrated Genomics Operation). We acknowledge Aaron Praiss, Ryan Kahn and Kaitlyn Gill (MSK) for their assistance with *in vivo* tumor harvest and/or processing.

## Funding

Research reported in this publication was supported in part by a Cancer Center Support Grant of the NIH/NCI (Grant No. P30CA008748), an Ovarian Cancer Research Alliance grant (#885167) and the Wasily Family Foundation. D.Z. is supported by the Ovarian Cancer Research Alliance Liz Tilberis Award, the Department of Defense Ovarian Cancer Research Academy (OC150111) and NCI R01 CA269382. B.W. is supported in part by Breast Cancer Research Foundation and Cycle for Survival grants. D.Z. and B.W are supported in part by BreakThrough Cancer Foundation grants. M.N.B. was supported by Fonds de recherche du Québec – Santé and the Ovarian Cancer Research Alliance.

## Author contributions

Conception and design: M.N.B., H.D., A.L.F.L., R.G., E.D., K.B.C., D.Z., B.W. Supervision: D.Z., B.W. Funding acquisition: M.N.B., D.Z., B.W. Bioinformatics analysis: Y.Z., A.G. Statistical analysis: M.N.B., M.G., Y.Z., A.G.,S.S., R.K. Experimental data acquisition: M.N.B., H.G., H.D., R.Q., Y.B., M.A.O., T.B., S.W. Data interpretation/analysis: M.N.B., H.D., Y.Z., A.L.F.L., A.G., S.S., R.K., P.J.H., Y.B., M.A.O., R.G., K.B.C., D.Z., B.W. M.N.B., D.Z. and B.W. drafted the manuscript, and M.N.B., T.B. and Y.B. prepared Figures. All authors reviewed the manuscript.

## Conflicts of interest

B.W. reports grant funding by Repare Therapeutics, outside the scope of the current study.

