## Supplementary material for "Interferon response and epigenetic modulation by *SMARCA4* mutations drive ovarian tumor immunogenicity": Table S1, Table S2, Table S3, Table S4, Table S5, Table S6, Table S7, Fig S1, Fig S2, Fig S3, Fig S4, Fig S5, Fig S6, Fig S7, Fig S8, Fig S9

Brodeur et al

Supplementary Tables S1 – S7

Supplementary Figures S1 – S9

**SUPPLEMENTARY TABLES**

**Table S1. Cell line characteristics**

| **Cell line** | **Derivation** | **Mutational Profile** | **Acquisition** |
|---|---|---|---|
| ID8-Luciferase | Mouse ovarian surface epithelium | *TP53*-, *ARID1A*-, *BRCA*-, *PIK3CA*-, *BRAF*-, *CTNNB1*- and *KRAS*-wildtype. Homologous-recombination proficient | Dr. Renier Brentjen's lab |
| UPK10-GFP | Traditional genetically engineered mouse model (GEMM) | *TP53*- and *KRAS*-mutated | Dr. Jose Conejo-Garcia lab |
| OAW28 | High-grade serous ovarian cancer patient | *TP53*- and *KRAS*-mutated | European Collection of Authenticated Cell Cultures |
| B16-F10 | Mouse melanoma | *BRAF*-wildtype, *TP53*-mutated | American Type Culture Collection |

**Table S2. Murine and human guide RNA sequences**

**Murine guide RNA (gRNA) sequences:**

| **SMARCA4 Exon** | **RNA sequence** |
|---|---|
| gRNA Exon 14 | CCCGACACATTATTGAGTAA |
| gRNA Exon 23 | TCACCGGCGGCATCGTGCAA |
| gRNA Exon 28 | TGCGGGCTTCTTCACGCCGG |

**Human guide RNA sequences**:

| ***SMARCA4* Exon** | **RNA sequence** |
|---|---|
| gRNA Exon 3 | TGGCCGAGGAGTTCCGCCCA |
| gRNA Exon 8 | TGGTTCGCAAATCCCCGGCC |
| gRNA exon 10 | TGGCCGAGGAGTTCCGCCCA |

**Table S3. Mouse and human primers for *SMARCA4* Sanger sequencing**

**Mouse primers**

| Exon | Forward | Reverse | Amplicon length (bp) |
|---|---|---|---|
| 14 | ATCATGGAGAAACGGTTTGG | GTAGGGGGCTGAGCCAAG | 543 |
| 23 | CTCTTCTGAGGCATCTCTTGC | CCGTATTTGGAGCCTTAGCA | 333 |
| 28 | GCTGGGATTACAGGCCAAT | CCGCAGTACCAGCTACACAT | 333 |

**Human primers**

| Exon | Forward | Reverse | Amplicon length (bp) |
|---|---|---|---|
| 3 | ATCATGGAGAAACGGTTTGG | GTAGGGGGCTGAGCCAAG | 543 |
| 8 | CTCTTCTGAGGCATCTCTTGC | CCGTATTTGGAGCCTTAGCA | 333 |
| 10 | GCTGGGATTACAGGCCAAT | CCGCAGTACCAGCTACACAT | 333 |

**Table S4. Short hairpin sequences**

| **Target gene** | **Short hairpin (sh) sequence** |
|---|---|
| shNTC | 5’TGCTGTTGACAGTGAGCGCTTAAATAACTACTGACGTCCGTAGTGAAGCCACAG ATGTACGGACGTC GTAGTTATTTAATTGCCTACTGCCTCGGA-3’ |
| shMAVS_1 | 5’-GCTGTTGACAGTGAGCGCGCATACATTGATGCTAATATATAGTGAAGCCACAG ATGTATATATTAGCATCAATGTATGCATGCCTACTGCCTCGGA-3’ |
| shMAVS_2 | 5’-CTGTTGACAGTGAGCGACCACACATACATGCTAATATATAGTGAAGCCACAGA TGTATATATTAGCATGTATGTGTGGGTGCCTACTGCCTCGGA-3’ |
| shIRF3 | 5’-TGCTGTTGACAGTGAGCGAGGAGGCTTAGCTGACAAAGAATAGTGAAGCCACA GATGTATTCTTTGTCAGCTAAGCCTCCGTGCCTACTGCCTCGGA-3’ |

**Table S5: Taqman probes and ERV primers**

### IFN genes

#### Taqman probes:

| Gene | Assay ID | Species |
|---|---|---|
| β-actin | Mm00607939_s1 | Murine |
| MDA5 | Mm00459183_m1 | Murine |
| RIG-1 | Mm00487934_m1 | Murine |
| IFNa4 | Mm00833969_s1 | Murine |
| IFNB | Mm00439552_s1 | Murine |
| STAT | Mm00439518_m1 | Murine |
| STING | Mm01158117_m1 | Murine |
| IRF7 | Mm00516793_g1 | Murine |
| cGAS | Mm01147496_m1 | Murine |
| OASL | Mm00496187_m1 | Murine |
| MX1 | Mm00487796_m1 | Murine |
| MAVS | Mm00523170_m1 | Murine |
| IRF3 | Mm00516784_m1 | Murine |

| | | |
|---|---|---|
| B-actin | Hs01060665_g1 | Human |
| IFI44L | Hs00199115_m1 | Human |
| IRF7 | Hs01014809_g1 | Human |
| STING | Hs00736955_g1 | Human |
| OASL | Hs00984387_m1 | Human |
| cGAS | Hs00403553_m1 | Human |
| IFNB1 | Hs01077958_s1 | Human |
| IFI27 | Hs01086370_m1 | Human |
| DDX41 | Hs00169602_m1 | Human |
| IFI44 | Hs00197427_m1 | Human |
| IFI16 | Hs00986757_m1 | Human |
| RIG-1 | Hs00204833_m1 | Human |
| MX-1 | Hs00895608_m1 | Human |
| STAT1 | Hs01013996_m1 | Human |

### ERV genes

#### Murine ERV or LINE primers:

| Gene | Forward | Reverse |
|---|---|---|
| B-actin | TTC TTG GGT ATG GAA TCC TGT GG | TGG CAT AGA GGT CTT TAC GGA TG |
| Syncytin-A | GAT GAC ATC CAC TGC CAC AC | ATT GTC CGG CTC GAA TAG G |
| mERVL-gag-pol | ACA TAC CCA GTA ATG GTC AGC AC | ATT GGT TAG CCA GTA CCA AAG GT |
| mERV3 | CAT AGC CTC TAC CTT CTG TCT GGT | AGA GGT CAT AGC ATT GTA GGG TTC |
| Peg11(Mart1)(Rtl1) | GAA ACA ATC AAC TCA TCC GAG AC | AGA GTT CTT GGG CTG ACC TTC |
| mMart8 (Cxx1c) | AAG GGC CGG GCC CTG CAG TG | CTA GAA GTC CTC ATC CTC CTC CCA CCC G |
| B1 (consensus) | GAG GCA GAG GCA GGC GGA TT | GTT TCT CTG TGT AGC CCT GGC |
| Mouse L1-Tf | CAG CGG TCG CCA TCT TG | CAC CCT CTC ACC TGT TCA GAC TAA |

#### Human ERV primers:

| Gene | Forward | Reverse |
|---|---|---|
| B-actin | CAA CCG CGA GAA GAT GAC C | TAG CAC AGC CTG GAT AGC AA |
| TBP | GAA GTT GGG TTT TCC AGC TAA GT | ACT AAA TTG TTG GTG GGT GAG C |
| ERV-Fc2 | CTC CAT TAG TAG CAG TTC CTC TCC | GAG AAT AGT GGG ACC TGT CCT TT |
| ERV-K | TTT GAC TGA AGT ATT AAA AGG TGT | GTG ACT GCA ATT AAT CCC ATA ATC |
| Syncytin-1 | ATG GAG CCC AAG ATG CAG | AGA TCG TGG GCT AGC AG |
| Syncytin-3 | TTA GCC ACA AAT TCT GGG ATA ACT | AGA GGT AAC AAT AGA GGC CAT GAG |

**Table S6. Flow cytometry antibodies**

**Extracellular antibodies used *in vivo* with ID8 model**

| Antibody | Catalog Number | Dilution |
|---|---|---|
| Zombie-NIR dye | 423106, BioLegend | 1:8000 |
| CD45-BUV395 | 564279, BD Biosciences | 1:40 |
| CD3-APC/Fire810 | 100267, Biolegend | 1:80 |
| CD4-BUV496 | 612952, BD Biosciences | 1:40 |
| CD8a-BUV805 | 752640, BD Biosciences | 1:20 |
| NK1.1-BUV661 | 741477, BD Biosciences | 1:40 |
| CD11b-BV605 | 101257, Biolegend | 1:40 |
| CD11c-BUV563 | 749040, BD Biosciences | 1:160 |
| F4/80-BUV737 | 749283, BD Biosciences | 1:40 |
| LY6G-PerCP | 127653, Biolegend | 1:40 |
| LY6C-AF700 | 128023, Biolegend | 1:100 |
| CD206-PE/Dazzle594 | 141732, Biolegend | 1:80 |
| B220-BV650 | 103241, Biolegend | 1:20 |
| I-A/I-E-BUV786 | 742894, BD Biosciences | 1:40 |
| CD86-APC | 17-0862-81, Invitrogen | 1:40 |
| PDL1-BV421 | 564716, BD Biosciences | 1:160 |
| ICOS-PE | 12-9942-81, Invitrogen | 1:80 |
| PD1-FITC | 11-9985-81, Invitrogen | 1:50 |

**Intracellular antibodies used *in vivo* with ID8 model**

| Antibody | Catalog Number | Dilution |
|---|---|---|
| FoxP3-PerCP-eF710 | 46-5773-82, Invitrogen | 1:320 |
| Granzyme B-PE-Cy7 | 25-8898-80, Invitrogen | 1:160 |

**Extracellular antibodies used *in vivo* with B16-F10 model**

| Antibody | Catalog Number | Dilution |
|---|---|---|
| Fixable Viability Dye eFluor506 | 65-0866-18, Invitrogen | 1:1000 |
| CD45.2-AF700 | 56-0454-82, Invitrogen | 1:400 |
| CD3-BUV496 | 364-0032-82, Invitrogen | 1:400 |
| CD11b-BV570 | 101233, BioLegend | 1:400 |
| CD11c-APC-Cy7 | 561241, BD Biosciences | 1:400 |
| F4/80-BV650 | 416-4801-82, Invitrogen | 1:400 |
| CD8a-Pacific Orange | MCD0830; Invitrogen | 1:400 |
| NK1.1-PerCP-Cy5.5 | 45-5941-82, Invitrogen | 1:400 |
| CD4-Qdot-605 | Q10092, Invitrogen | 1:400 |
| MHCII-eFluor450 | 48-5321-82, Invitrogen | 1:400 |

**Intracellular antibodies used *in vivo* with B16-F10 model**

| Antibody | Catalog Number | Dilution |
|---|---|---|
| FoxP3/APC | 17-5773, Invitrogen | 1:200 |

**Extracellular antibodies used in vitro with OAW28 and ID8 models**

| Antibody | Catalog Number | Dilution |
|---|---|---|
| Viability Zombie-Green dye | 423111, BioLegend | 1:2000-4000 |
| Anti-mouse PDL1-APC | 564715, BD Biosciences | 1:40 |
| Anti-mouse H-2Kb-PE/Cy7 | 116519, BioLegend | 1:160 |
| Anti-human PDL1-APC | 568316, BD Biosciences | 1:80 |
| Anti-human HLA-ABC-PE/Cy7 | 25-9983-42, eBioscience | 1:640 |

**Table S7: *SMARCA4* RNA expression and *SMARCA4* mutation type from The Cancer Genome Atlas pan-cancer study**

| **Mutation type** | ***SMARCA4*-low expression** (n) | ***SMARCA4*-high expression** (n) | **p-value** |
|---|---|---|---|
| *SMARCA4* wild-type | 1914 | 2008 | - |
| *SMARCA4* truncating/ splice mutation | 45 | 3 | P<0.0001 |
| *SMARCA4* missense mutation | 43 | 67 | P=0.0527 |
| *SMARCA4* UTR mutation | 1 | 0 | P=0.4881 |

* P-values were calculated using the Fisher exact test. P-value <0.05 was considered statistically significant.

### SUPPLEMENTARY FIGURES

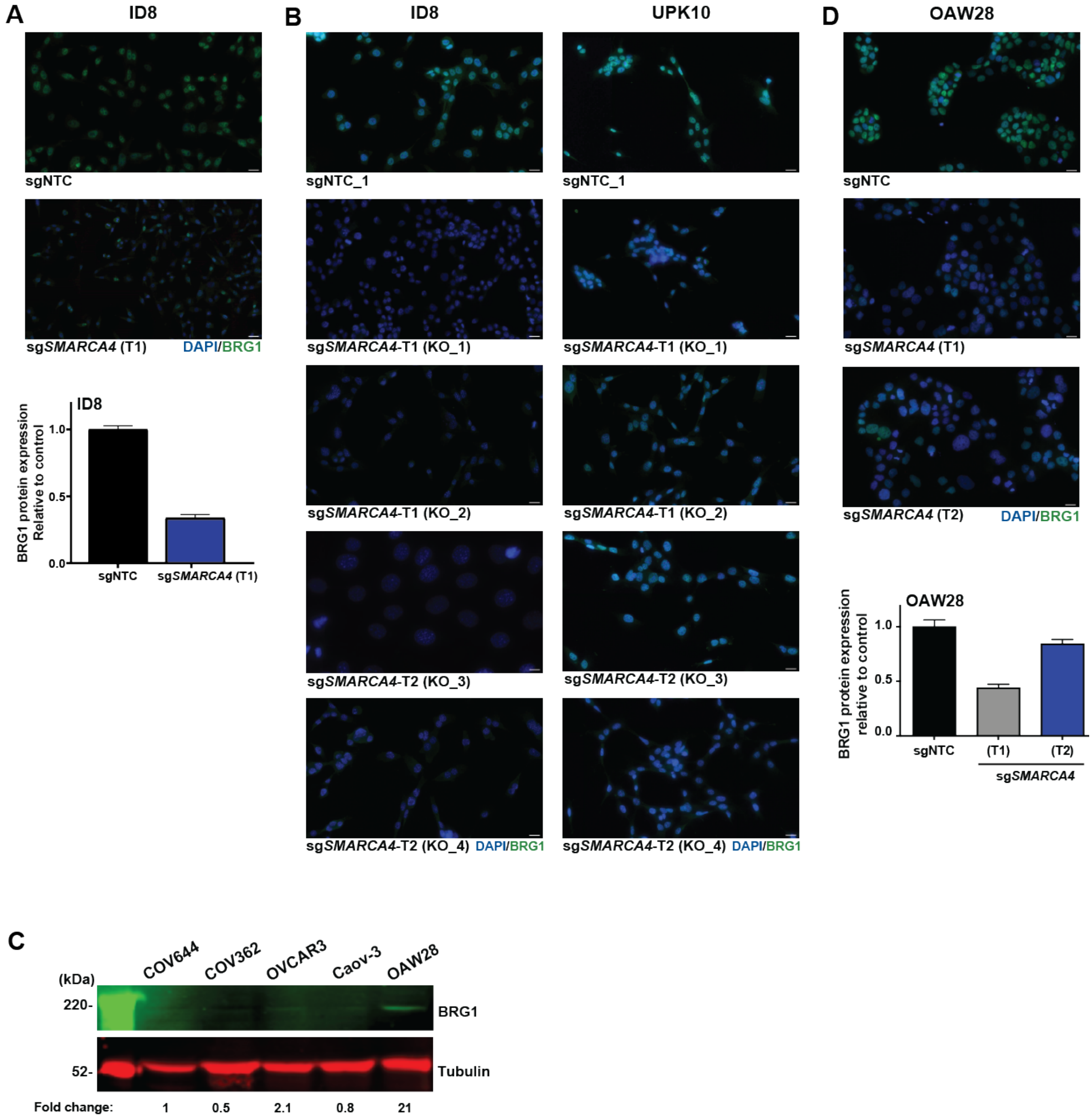


**Figure S1. Development of single guide (sg)SMARCA4 and sgNTC ovarian cancer models**

**A)** ID8 non-clonally selected models (sgNTC + sg*SMARCA4*) with BRG1 protein quantification. **B)** ID8 and UPK10 clones for sgNTC (1 clone) and sg*SMARCA4* (4 clones) models. **C)** BRG1 protein expression in a panel of 5 human OC cell lines. **D)** OAW28 non-clonally selected models (sgNTC + sg*SMARCA4*) with BRG1 protein quantification.

*P<0.05, **P<0.01, ***P<0.001, ****P<0.0001. Error bars represent ± SEM. ID8/UPK10-T1 = target exon 14, ID8/UPK10-T2 = target exon 23, KO_1 = target exon 14 clone 1, KO_2 = target exon 14 clone 2, KO_3 = target exon 23 clone 3, KO_4 = target exon 23 clone 4, NTC = non-target control, NTC_1 = NTC clone 1, OAW28-T1 = target exon 3, OAW28-T2 = target exon 8.

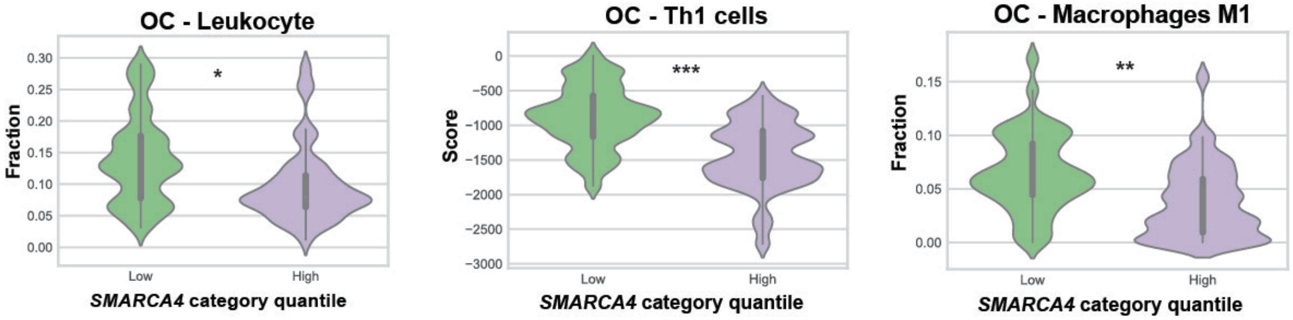


**Figure S2.** ***SMARCA4*** **expression in ovarian cancers with immune related parameters.**
Immune signatures and cell populations by *SMARCA4* expression derived from RNA sequencing data for high-grade serous ovarian cancers obtained from The Cancer Genome Atlas.
Statistical analysis was performed using Fisher exact test and Benjamini-Hochberg adjustment. *P<0.05, **P<0.01, ***P<0.001, ****P<0.0001.

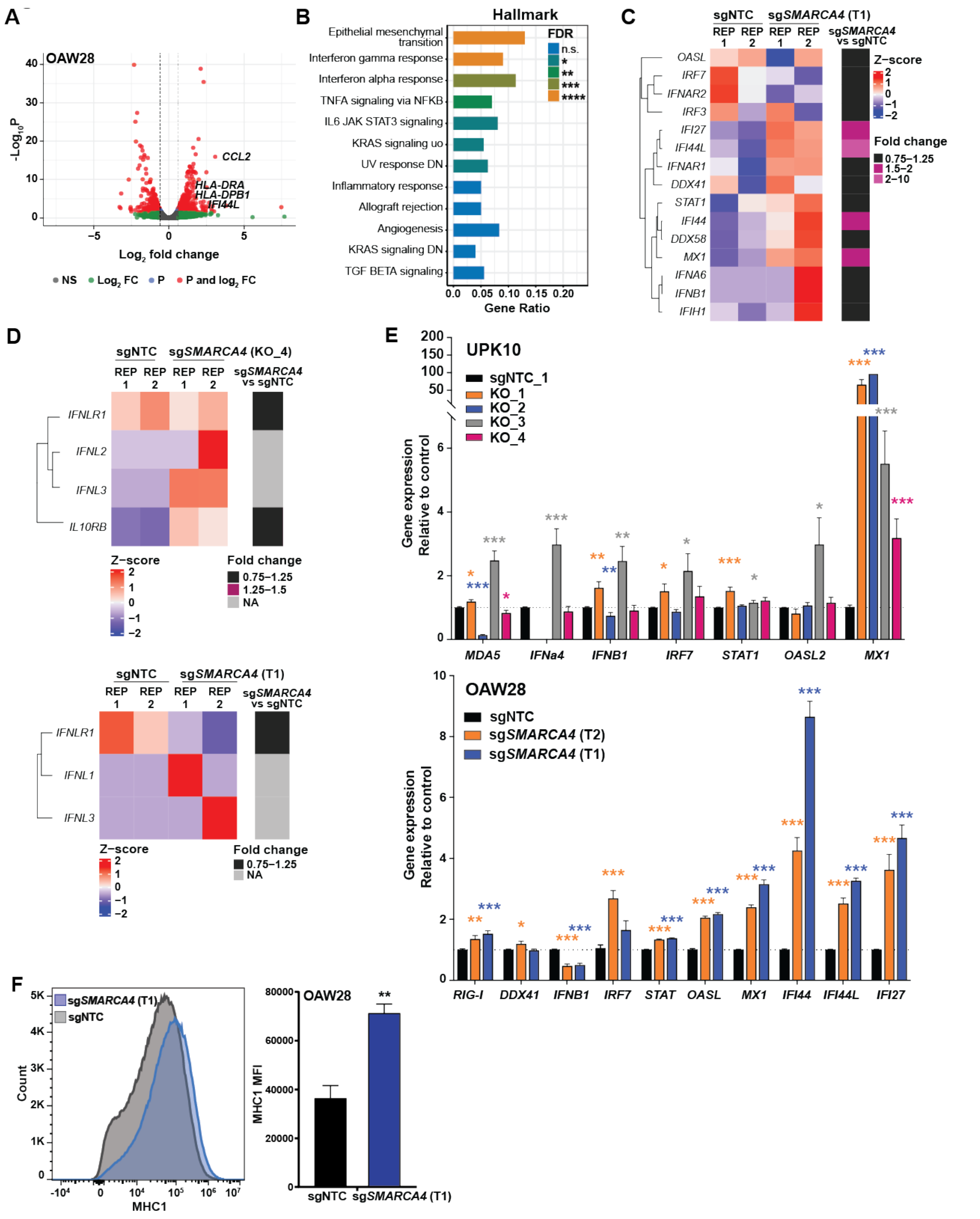


**Figure S3. Loss of function mutations of *SMARCA4* in ovarian cancer cells leads to increased interferon response and antigen presentation gene activation.**

**A)** Volcano plot for differential expression of genes in sg*SMARCA4* compared to sgNTC OAW28 cells. Specific genes are labeled. **B)** Upregulated Hallmark pathways in sg*SMARCA4* compared to sgNTC

OAW28 cells. Statistical analysis was based on hypergeometric test and performed using ClusterProfiler. **C)** Gene expression heatmap of type I IFN genes in OAW28 cells. RPKM values were scaled to Z-score for visualization. Gene expression fold-change is color-coded according to the legend. **D)** Gene expression heatmap of type III IFN genes in ID8 (top) and OAW28 (bottom) cells. Gene expression fold-change is color-coded according to the legend. **E)** qRT-PCR validation results for IFN genes using UPK10 (sgNTC + 4 sg*SMARCA4* clones) and OAW28 (sgNTC + 2 sg*SMARCA4* targets). Expression levels were normalized to B-actin expression, and comparisons of mRNA expression levels were performed relative to control (sgNTC). **F)** MHC1 expression in ID8 cells by flow cytometry with quantification. Statistical analysis was performed using two-tailored unpaired t-test (**E, F**). *P<0.05, **P<0.01, ***P<0.001, ****P<0.0001. Error bars represent ± SEM. Samples in duplicates (**A, B, C, D**) and triplicates (**F**). N=3 independent experiments (**E**). KO_1 = target exon 14 clone 1, KO_2 = target exon 14 clone 2, KO_3 = target exon 23 clone 3, KO_4 = target exon 23 clone 4, NTC = non-target control, NTC_1 = NTC clone 1, OAW28-T1 = target exon 3, OAW28-T2 = target exon 8.

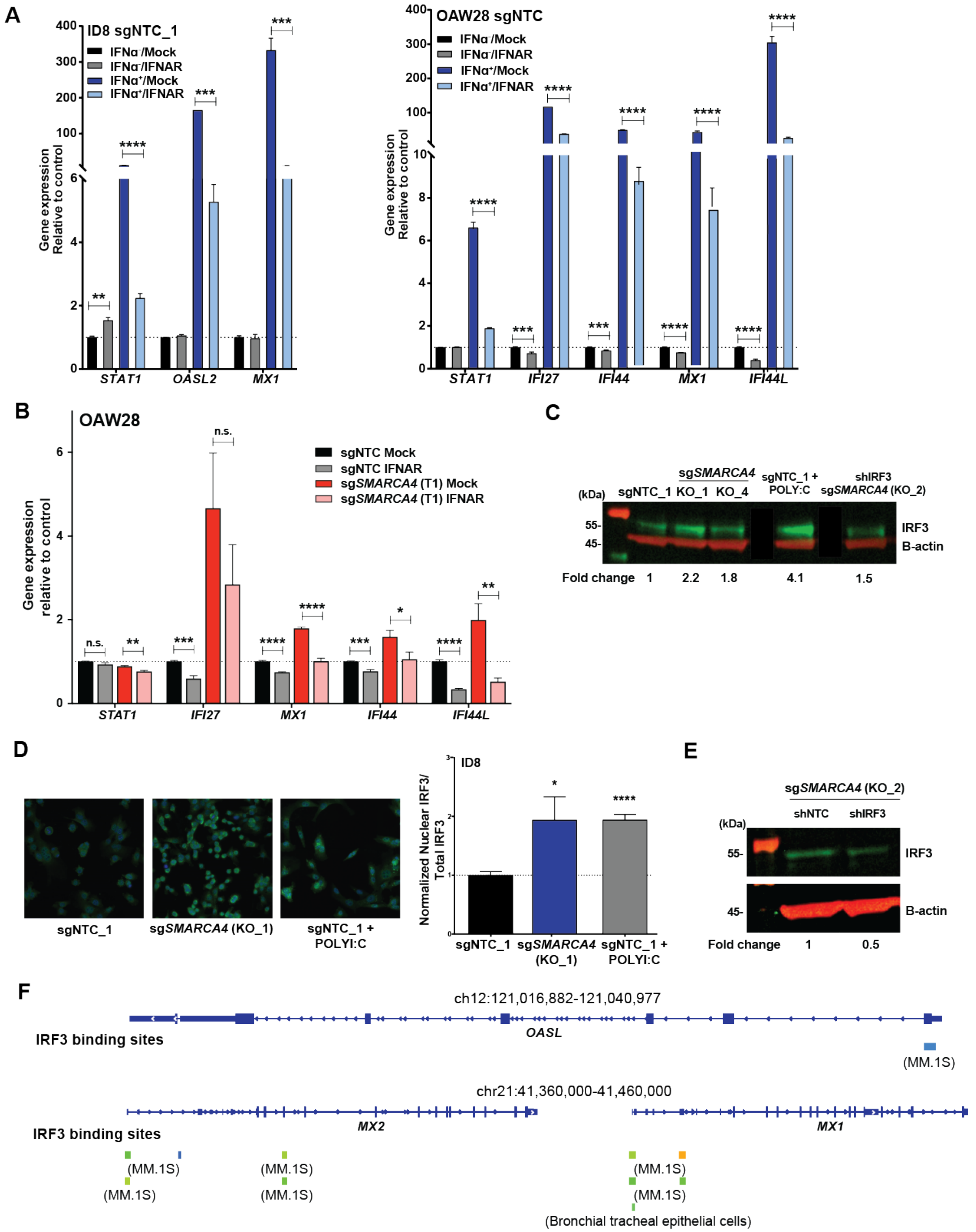


**Figure S4. *SMARCA4* loss of function upregulates immune genes through type I interferon signaling.**

**A)** Type I IFN receptor (IFNAR) neutralizing assay for dose-effectiveness in ID8 and OAW28 cells using IFNα (or control media) at 2000U/mL for 24 hours followed by IFNAR antibody at 10 and 8 ug/mL, respectively, for 48 hours. **B)** IFNAR-2 neutralizing assay in OAW28 sgNTC versus sg*SMARCA4* cells. **C)** IRF3 protein expression in sg*SMARCA4* vs sgNTC cells (POLYI:C for positive control and shIRF3 for negative control). B-actin was used as protein loading control. Relative quantification (below) of protein levels as compared to control. **D)** Activated IRF3 protein levels in sg*SMARCA4* vs sgNTC cells (POLYI:C for positive control). **E)** IRF3 protein expression in shNTC- and shIRF3-transfected sg*SMARCA4* cells by western blot. B-actin was used as protein loading control. Relative quantification (below) of protein levels as compared to control. **F)** Analysis of publicly available ChIP-Atlas data of IRF3 DNA binding sites on ISGs in human cancer cell lines.

For qRT-PCR experiments (**A, B**), expression levels were normalized to B-actin expression, and comparisons of mRNA expression levels were performed relative to control (sgNTC). Statistical analysis was performed using two-tailored unpaired t-test (**A, B, D**). *P<0.05, **P<0.01, ***P<0.001, ****P<0.0001. Error bars represent ± SEM. N=3 independent experiments (**A, B**). KO_1 = target exon 14 clone 1, KO_2 = target exon 14 clone 2, KO_4 = target exon 23 clone 4, NTC = non-target control, NTC_1 = NTC clone 1, OAW28-T1 = target exon 3, POLYI:C = Polyinosinic-polycytidylic acid.

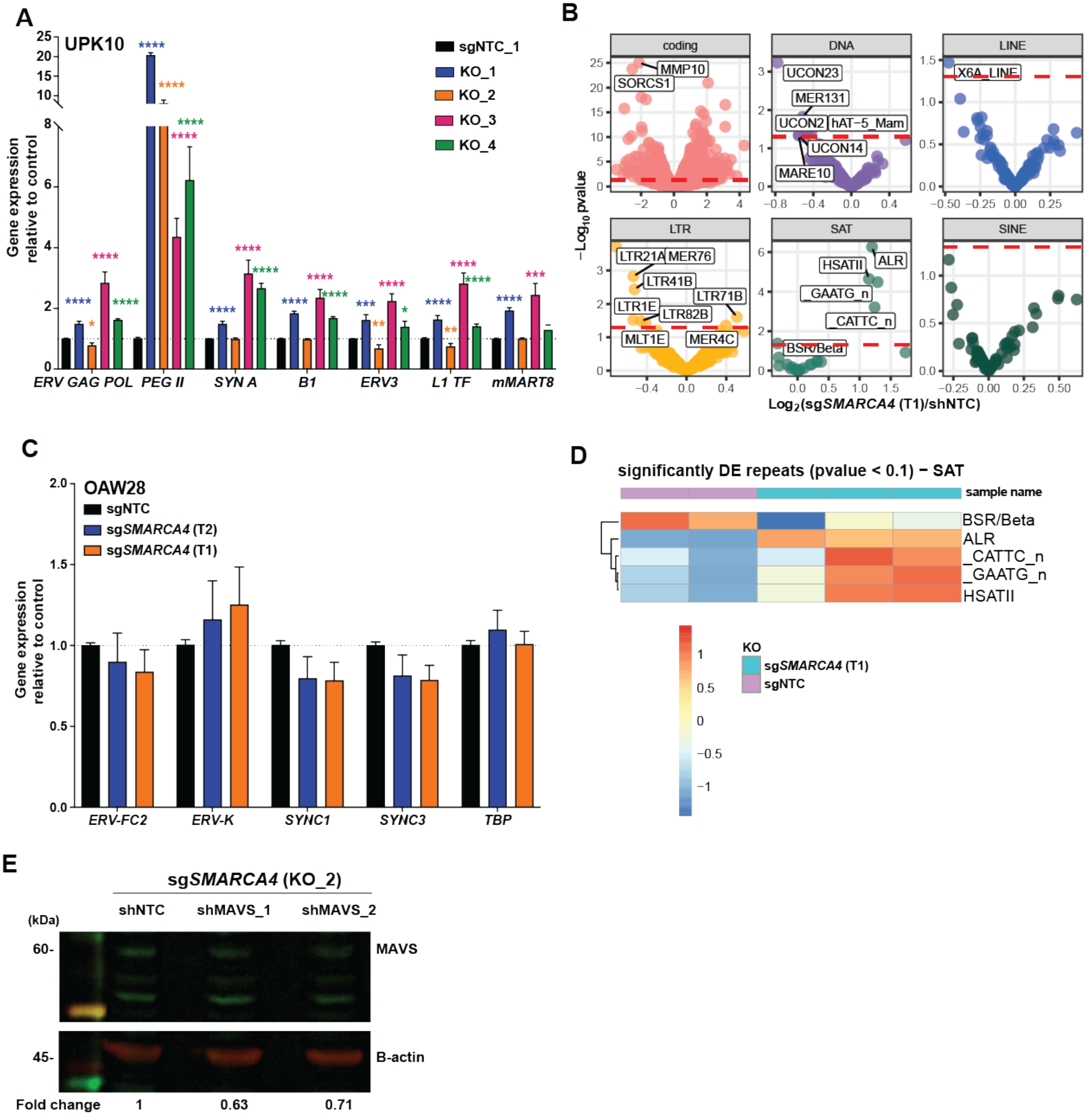


**Figure S5. Increase in ISGs is mediated through the dsRNA sensing pathway.**

**A)** qRT-PCR quantification of a panel of murine ERV genes in UPK10 sgNTC versus sg*SMARCA4* clones. **B)** Volcano plots for differential expression of TEs in OAW28 cells. **C)** qRT-PCR quantification of a panel of human ERV genes in OAW28 sgNTC and sg*SMARCA4* cells. **D)** Heatmap of SAT repeats with p<0.1. **E)** MAVS expression in shNTC- and shMAVS-transfected sg*SMARCA4* cells by western blot. B-actin was used as protein loading control. Relative quantification (below) of protein levels as compared to control.

For qRT-PCR experiments (**A, C**), expression levels were normalized to B-actin expression, and comparisons of mRNA expression levels were performed relative to control (sgNTC). Red line in (**B**) represents a cut-off of an adjusted p-value <0.05. *P<0.05, **P<0.01, ***P<0.001, ****P<0.0001. Error bars represent ± SEM. N=3 independent experiments (**A, C**). Samples in at least duplicates (**B, D**). KO_1 = target exon 14 clone 1, KO_2 = target exon 14 clone 2, KO_3 = target exon 23 clone 3, KO_4 = target exon 23 clone 4, NTC = non-target control, NTC_1 = NTC clone 1, OAW28-T1 = target exon 3, OAW28-T2 = target exon 8.

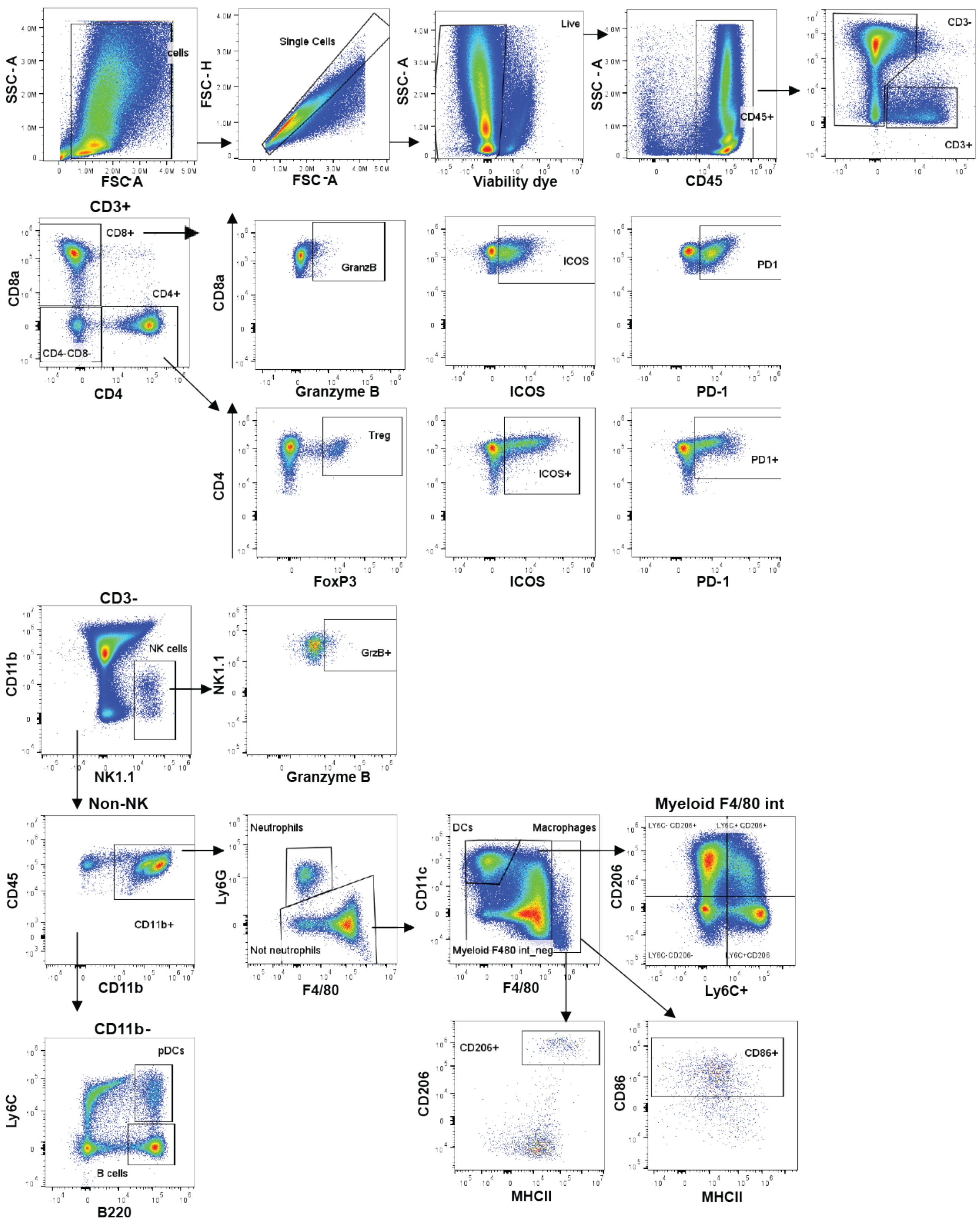


**Figure S6.** Gating strategy for spectral flow cytometry multi-parameter analysis using FlowJo in ID8 tumors (20-marker panel)

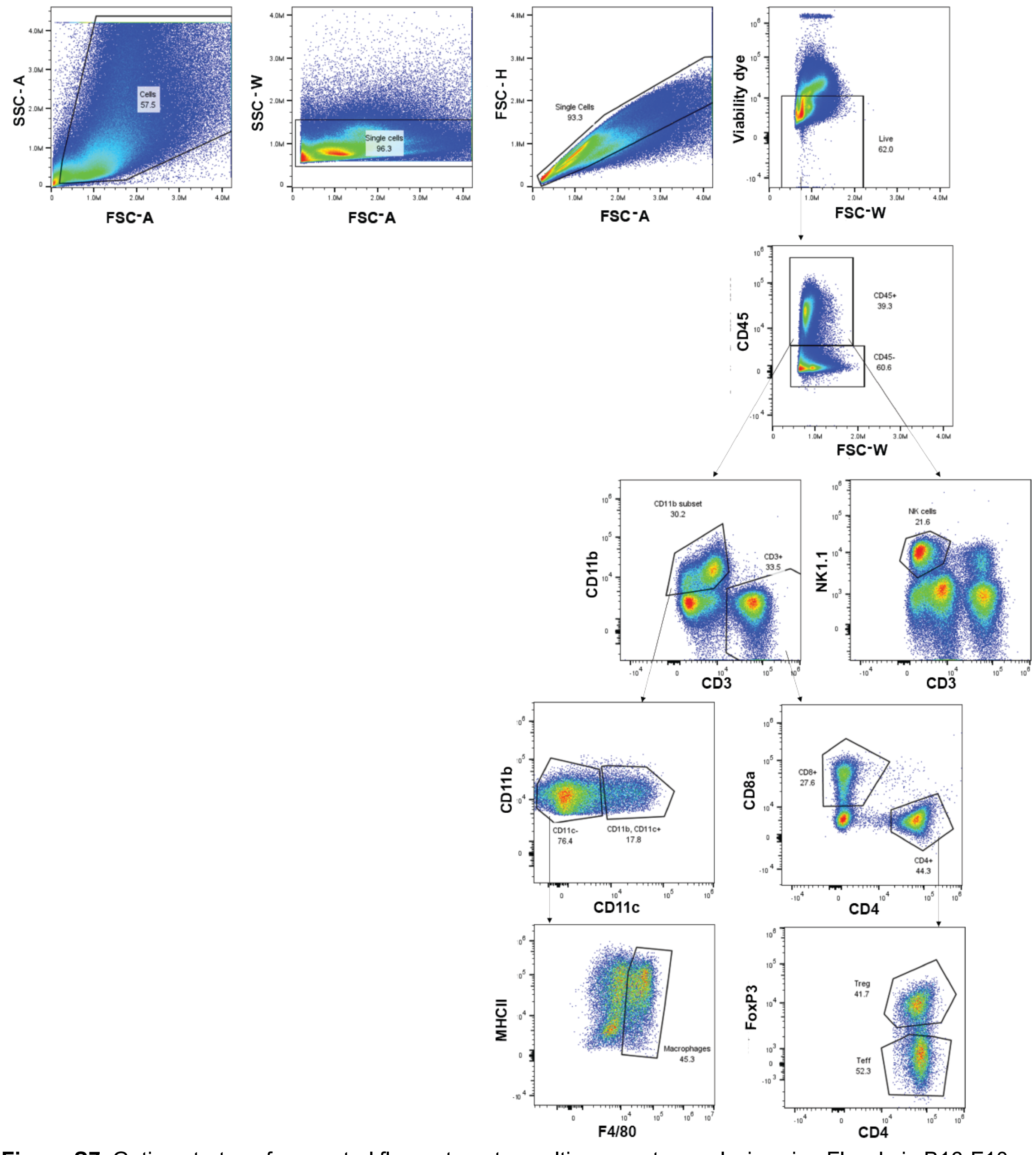


**Figure S7.** Gating strategy for spectral flow cytometry multi-parameter analysis using FlowJo in B16-F10 tumors (16-marker panel)

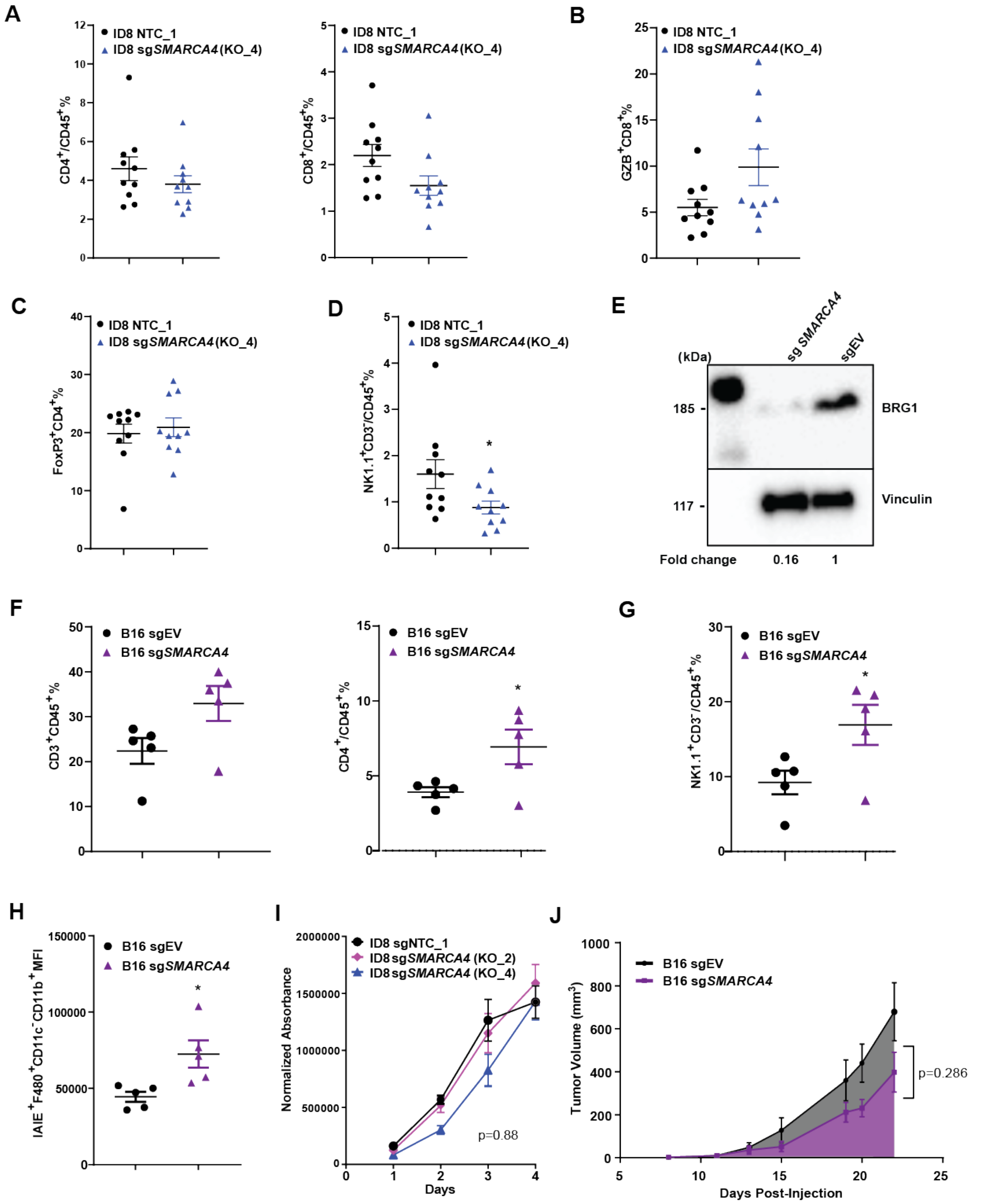


**Figure S8. *SMARCA4* loss of function leads to increase in vivo immunogenicity.**
**A)** Frequency of tumor CD4+ T cells (left) and CD8+ T cells (right) in ID8 model (of CD45+ cells). **B)** Frequency in Granzyme B expressing CD8+ T cells in ID8 model. **C)** Frequency of tumor T regulatory cells (of CD45+ cells). **D)** Frequency of tumor NK1.1+ cells in ID8 model (of CD45+ cells). **E)** Western

blot of sg*SMARCA4* and sgEV B16 melanoma model. Vinculin was used as protein loading control. Relative quantification (below) of protein levels as compared to control. **F)** Frequency of tumor CD3+ cells (left) and CD4+ T cells (right) in B16 model (of CD45+ cells). **G)** Frequency of NK1.1+ cells in B16 model (of CD45+ cells). **H)** MFI of MHCII expressing macrophages. **I)** Cell titer blue proliferation assay of ID8 sgNTC vs sg*SMARCA4* cells. N=3 independent experiments. Two-way ANOVA analysis was performed with exact p-value indicated on panel. **J)** Tumor volume in sg*SMARCA4* vs sgNTC B16 tumors (unpaired t-test of area under the curve with p-value indicated on panel). Statistical analysis was performed using two-tailored unpaired t-test (**A, B, C, D, F, G, H**). *P<0.05, **P<0.01, ***P<0.001, ****P<0.0001. Error bars represent ± SEM. N=10 mice/group in **A, B, C, D**, and N=5 mice/group in **F, G, H, J**. EV = empty vector, KO_2 = target exon 14 clone 2, KO_4 = target exon 23 clone 4, NTC = non-target control, NTC_1 = NTC clone 1.

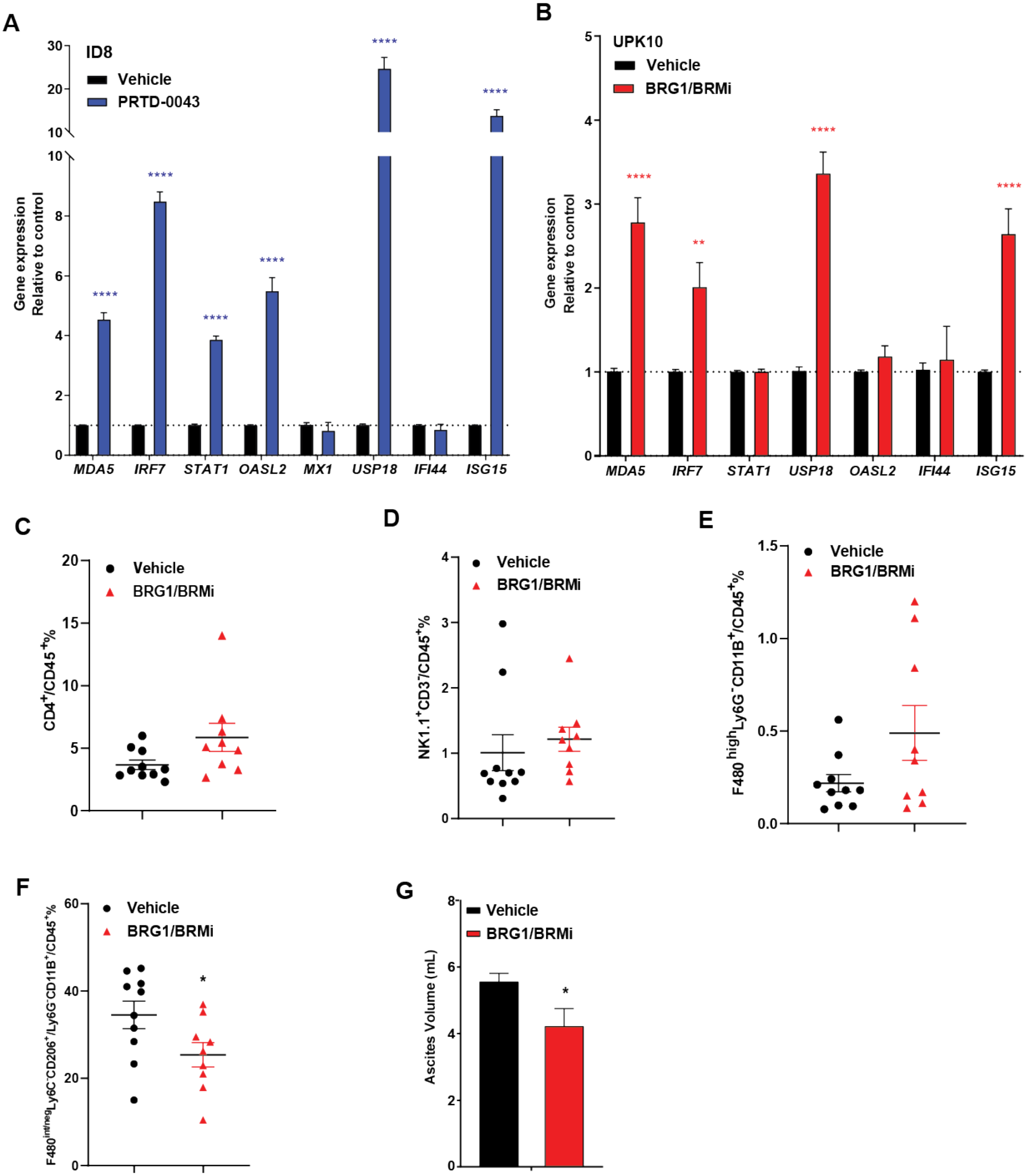


**Figure S9. Therapeutic targeting of BRG1 recapitulates the immunogenic effects of *SMARCA4* loss.**

**A)** qRT-PCR quantification of IFN gene expression using ID8 parental cell treated with BRG1/BRM protein degrader or vehicle **B)** qRT-PCR quantification of IFN gene expression using UPK10 parental cell treated with BRG1/BRM inhibitor or vehicle. **C)** Frequency of tumor CD4+ T cells (of CD45+ cells). **D)** Frequency of tumor NK1.1+ cells (of CD45+ cells). **E)** Frequency of tumor dendritic cells (of CD45+ cells). **F)** Frequency of tumor macrophages (of CD45+ cells). **G)** Ascites volume (tumor burden) measured at week 3.

For qRT-PCR experiments (**A, B**), expression levels were normalized to B-actin expression, and comparisons of mRNA expression levels were performed relative to control (vehicle). Statistical analysis was performed using two-tailored unpaired t-test. *P<0.05, **P<0.01, ***P<0.001, ****P<0.0001. Error bars represent ± SEM. N=10 mice/group in **C, D, E, F, G**.
